# Global analysis of methionine oxidation provides a census of folding stabilities for the human proteome

**DOI:** 10.1101/467290

**Authors:** Ethan J. Walker, John Q. Bettinger, Kevin A. Welle, Jennifer R. Hryhorenko, Sina Ghaemmaghami

## Abstract

The stability of proteins influences their tendency to aggregate, undergo degradation or become modified in cells. Despite their significance to understanding protein folding and function, quantitative analyses of thermodynamic stabilities have been mostly limited to soluble proteins in purified systems. We have used a highly multiplexed proteomics approach, based on analyses of methionine oxidation rates, to quantify stabilities of ~10,000 unique regions within ~3,000 proteins in human cell extracts. The data identify lysosomal and extracellular proteins as the most stable ontological subsets of the proteome. We show that the stability of proteins impacts their tendency to become oxidized and is globally altered by the osmolyte trimethylamine N-oxide (TMAO). We also show that most proteins designated as intrinsically disordered retain their unfolded structure in the complex environment of the cell. Together, the data provide a census of the stability of the human proteome and validate a methodology for global quantitation of folding thermodynamics.

**SIGNIFICANCE STATEMENT:** Most natural proteins fold into a native folded conformation stabilized by non-covalent interactions. The free energy difference between the folded and unfolded conformations of a protein (‘thermodynamic folding stability’) establishes the fraction of its population that is in a folded conformation at equilibrium. Here, we used a mass spectrometry-based approach to measure the stabilities of ~10,000 unique domains within ~3,000 proteins in human cell extracts. Our data provide a global census of the magnitudes of folding stabilities within the human proteome and identifies specific subsets of the proteome that are enriched in stable and unstable domains. We also show that the stability of proteins impacts their tendency to become oxidized and is globally altered by the chemical chaperone trimethylamine N-oxide (TMAO).

## INTRODUCTION

Within a cell, most proteins exist in a dynamic equilibrium between folded and unfolded conformations (1, 2). The free energy difference between these two states (ΔG_folding_ or ‘thermodynamic folding stability’) establishes the fraction of its population that is in a folded conformation at equilibrium (3). Folding stabilities can impact the tendency of proteins to aggregate, oxidize or undergo degradation, and alterations in protein stabilities play a role in diverse biological processes and physiological disorders (e.g. aging and neurodegenerative proteinopathies) (4–8). Despite their fundamental importance to protein function, the magnitude and span of folding stabilities within the proteome remain unclear. For example, estimations of the fraction of the proteome that harbors ‘intrinsically disordered’ domains are largely based on theoretical predictions and, with a few exceptions (9, 10), have yet to be verified by proteome-wide experimental data (11).

Historically, quantitative measurements of protein folding stabilities have been conducted for proteins in purified systems using *in vitro* denaturation experiments that employ global folding probes such as circular dichroism or isothermal calorimetry (12). Although these studies have made significant contributions to our understanding of folding thermodynamics, they have inherent limitations that mitigate their *in vivo* relevance. First, these studies have been biased towards well-folded stable protein model systems that can be easily expressed and purified. Hence, the thermodynamic folding parameters measured for well-studied folding models may not be representative of the proteome as a whole. Second, experiments conducted in purified systems typically do not account for the effects of interacting partners and other cellular factors that may influence protein stabilities *in vivo.* Third, these methodologies are inherently low throughput and cannot be easily used to investigate protein stabilities on proteome-wide scales.

To address these shortcomings, a number of recently developed techniques have investigated protein folding patterns using probes that can be analyzed by mass spectrometry-based proteomics. Hydrogen/deuterium exchange (HX) (13, 14), hydroxyl radical footprinting (15) and limited proteolysis (9, 16–19) have been coupled with liquid chromatography-tandem mass spectrometry (LC-MS/MS) to provide proteome-wide measures of protein structure. These global studies have successfully investigated thermostabilities (melting temperatures), ligand binding interactions and conformational changes induced by environmental alterations. However, due in part to technical limitations inherent in each of these approaches, it has proven difficult to obtain measurements of thermodynamic folding stabilities (ΔG_folding_) on proteome-wide scales.

In this study, we globally measured protein folding stabilities by using methionine oxidation as a probe. This approach, referred to as Stability of Proteins from Rates of Oxidation (SPROX) was initially described by West and colleagues (20) and is conceptually similar to previously described probing methodologies based on thiol labeling (21). SPROX offers a number of advantages that are particularly conducive to proteomic workflows. Methionine oxidation is a small, non-labile modification that can be readily coupled with chemical denaturation and is retained during the course of LC-MS/MS analyses. Selective oxidation of methionines allows for straightforward database searches, identification of modification sites and interpretation of oxidation kinetics.

Furthermore, the use of chemical denaturants prevents the irreversible aggregation of proteins during the course of denaturation – a phenomenon that typically confounds thermal unfolding experiments. To date, SPROX has been used to analyze the stability of individual intact proteins and to examine proteome-wide alterations in denaturation patterns induced by ligand binding (20, 22, 23). Here, we have coupled SPROX with highly multiplexed tandem mass tagging (TMT) to obtain high-resolution denaturation curves – an approach we have termed high resolution SPROX (HR-SPROX). By analyzing HR-SPROX data in the context of thermodynamic protein folding models, we were able to measure localized ΔG_folding_ values on a proteome-wide scale.

## RESULTS

### HR-SPROX workflow

Figure 1 outlines the premise and workflow of HR-SPROX analyses and detailed methods are provided in Supplementary Information. Methionine is a readily oxidizable amino acid and can be converted to methionine sulfoxide by the oxidizing agent hydrogen peroxide (H_2_O_2_) (24). The oxidation rate of a methionine residue within a protein is contingent on its local structural environment. Exposed methionines in unfolded regions oxidize faster than protected methionines in folded regions (25–27). Thus, the energy difference between the folded and unfolded conformations (ΔG_folding_) can modulate the folding equilibrium and rate of methionine oxidation. The progressive addition of a chemical denaturant results in the unfolding of the protein (as ΔG_folding_ increases) and escalates the rate of oxidation. Monitoring the extent of oxidation (at a constant oxidation time) in the presence of increasing concentrations of a chemical denaturant produces a denaturation curve (20).

**Figure 1.**
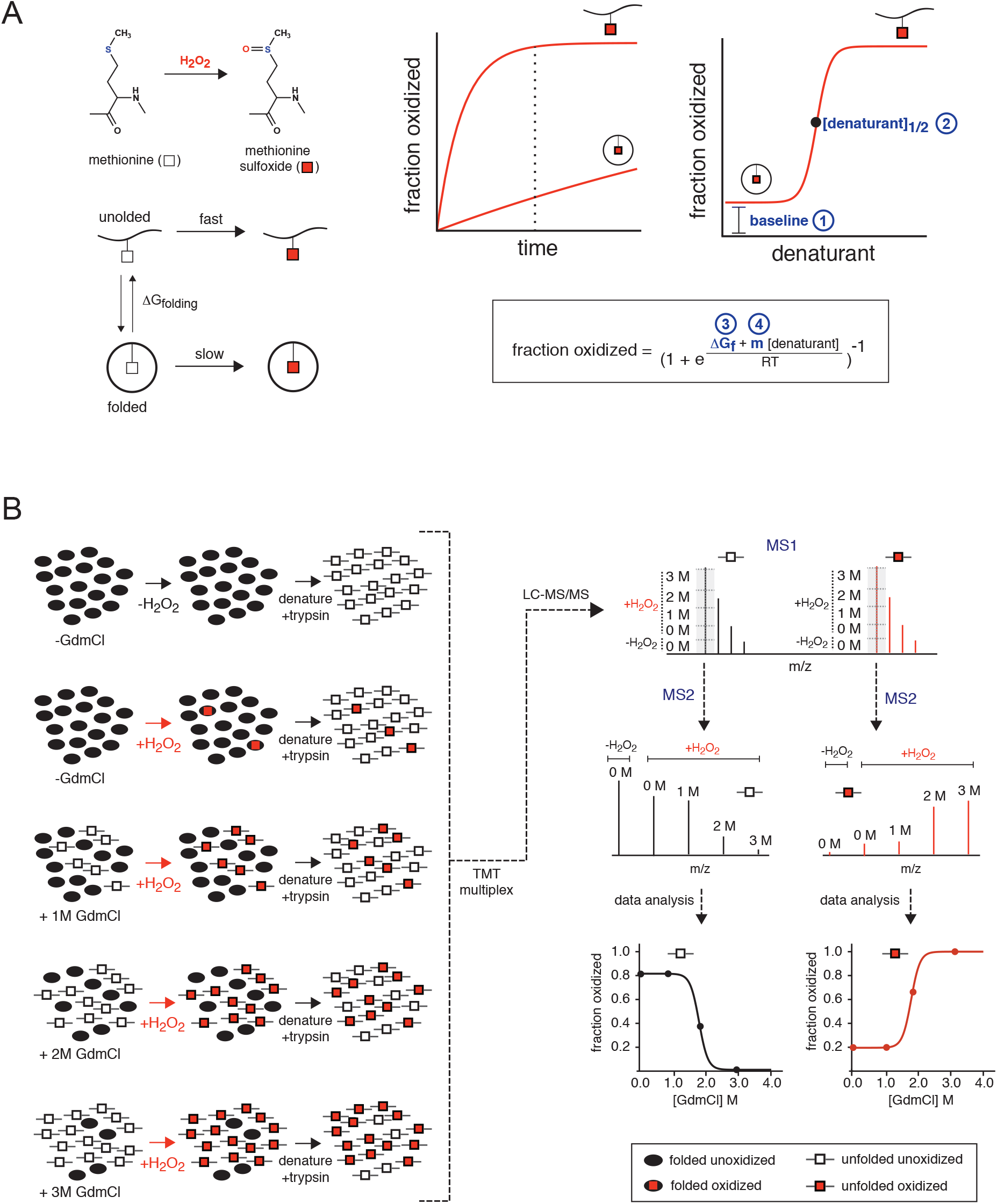
HR-SPROX as a tool for measuring protein stability. (A) Theoretical basis of SPROX. Methionine residues (white squares) are oxidized by H_2_O_2_ to form methionine sulfoxides (red squares). Methionine oxidation rates differ between folded and unfolded states. Thus, protein denaturation can be quantified by measuring the extent of oxidation across different denaturant concentrations at a constant oxidation time (dashed line). The resulting denaturation curves contain 4 categories of information (numbered 1-4): baseline oxidation, midpoint of denaturation ([denaturant]_1/2_), m-value, and free energy of folding (ΔG_folding_ or ΔG_f_). For two-state proteins, the latter two parameters can be determined by fitting the denaturation curve to an equation derived from the linear extrapolation model (LEM). (B) HR-SPROX workflow. Cells are lysed under native conditions and folded proteins (black ovals) are unfolded with increasing concentrations of GdmCl. Methionines are converted to methionine sulfoxides (red squares) by addition of H_2_O_2_. A control experiment lacking H_2_O_2_ is included and used as a normalization point. Extracts are digested into peptides and each sample (corresponding to a different denaturant concentration) is labeled with a unique tandem mass tag (TMT) and subsequently combined and analyzed by LC-MS/MS. Reporter ion intensities at the MS2 level are internally normalized to create denaturation curves, monitoring either the increase in methionine sulfoxide-containing peptides or the decrease in unoxidized methionine-containing peptides.

The SPROX denaturation curve can potentially provide four types of information: baseline oxidation, [denaturant]_1/2_, ΔG_folding_ and m-value (Figure 1A). The level of baseline oxidation (the extent of oxidation induced by H_2_O_2_ in the absence of denaturant) provides a measure of the susceptibility of the methionine to oxidation in the context of the native protein. As we show below, this parameter is largely influenced by the level of solvent exposure of the methionine sidechain. The midpoint of denaturation ([denaturant]_1/2_) is a model-independent parameter that provides the denaturant concentration required to unfold half of the protein population. For proteins that conform to a two-state folding model, where only the fully folded and fully unfolded conformations are significantly populated at all denaturant concentrations, the resulting sigmoidal denaturation curve can be interpreted in terms of the energy difference between the two conformations using the following equation (see Supplementary Information for formal derivation and assumptions):

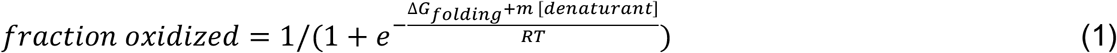

Where ΔG_folding_ is the folding stability in the absence of denaturant, and the m-value is the slope of the linear relationship between ΔG_folding_ and [denaturant] in accordance to the linear extrapolation model of two-state folding (LEM) (28). It has been shown that the m-value is correlated with the size and accessible surface area of the folding unit (29). Thus, for proteins or domains that unfold in a two-state fashion, least-squares fitting the SPROX curve with the above equation can provide ΔG_folding_ measurements for the folding unit that encompasses the methionine.

Accurate determination of ΔG_folding_ values from HR-SPROX curves using LEM is contingent on the availability of a significant number of data points in the transition regions of the sigmoidal curves (28). To obtain sufficient resolution required for analysis of a large number of polypeptides with diverse stabilities, we obtained data points at numerous denaturation concentrations (see below for exact statistics) and multiplexed the samples using tandem mass tags (TMT) (Figure 1B). Each methionine can potentially produce two HR-SPROX curves based on the disappearance of methionine or appearance of methionine sulfoxide as a function of denaturant concentration. For a given methionine, these two curves are expected to be mirror images of one another, providing two independent datasets for analysis of its localized stability.

It should be noted that methionine residues have a tendency to become oxidized in the cell or during mass spectrometric analysis, even in the absence of H_2_O_2_. However, the level of such spurious oxidation is minimal relative to the forced oxidation induced by H_2_O_2_. Furthermore, this background oxidation is expected to be constant at all denaturant concentrations and does not influence the measurements of normalized fractional oxidation.

### Analysis of purified lysozyme validates the accuracy of ΔG_folding_ measurements by HR-SPROX

We first validated the HR-SPROX approach by analyzing purified lysozyme, a well-studied two-state protein folding model. Lysozyme has three buried tryptophan residues whose fluorescence emission spectra change upon denaturation (Figure 2A). A protected methionine resides in the same region of the protein, enabling parallel analysis of stability by HR-SPROX. We obtained fluorescence and HR-SPROX denaturation curves using guanidine hydrochloride (GdmCl) as a denaturant. The experiments were carried out at room temperature at pH 7.4. Figure 2B compares the denaturation curves obtained by fluorescence and HR-SPROX for a methionine-containing tryptic peptide that also encompasses two of the buried tryptophan residues. Analysis of the two denaturation curves with the two-state folding model provides similar measurements for [GdmCl]_1/2_, m-value and ΔG_folding_. These values are also consistent with previous stability measurements obtained for lysozyme under similar conditions (30).

**Figure 2.**
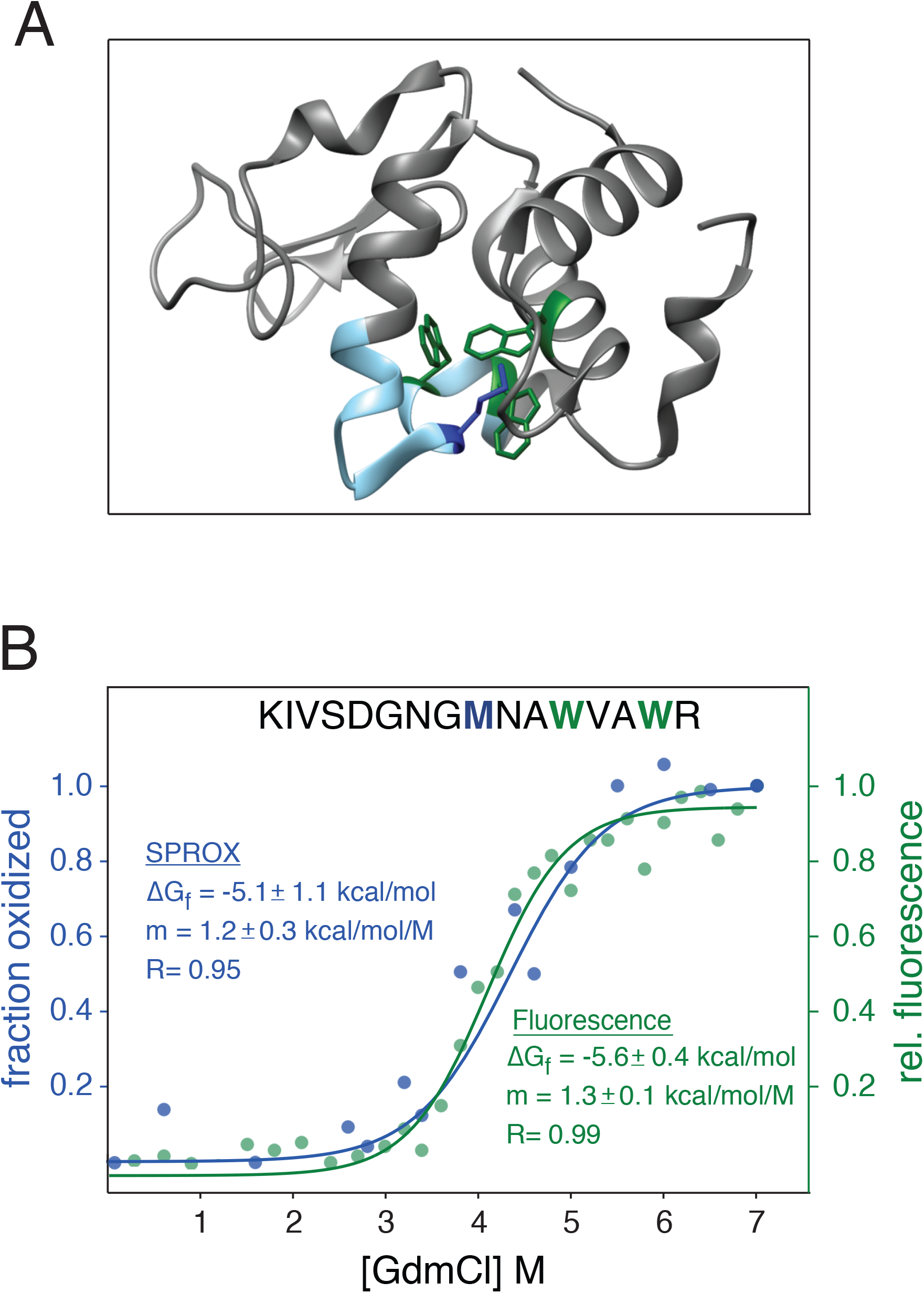
Validation of HR-SPROX by analysis of purified lysozyme. (A) The structure of hen lysozyme (1DPX). Buried tryptophan residues analyzed by fluorescence are colored in green, and the buried methionine residue analyzed by SPROX is colored in blue. (B) Denaturation curves obtained by intrinsic fluorescence (green) and SPROX (blue). The data were fit with Equation 1 to determine ΔG_folding_ and m-values.

### HR-SPROX enables proteome-wide analysis of folding thermodynamics in human fibroblast extracts

We next extended our analysis to the proteome-wide investigation of complex extracts. Human diploid fibroblasts expressing the catalytic component of human telomerase (HCA2-hTert) (31) were grown to a contact-inhibited quiescent state in culture and extracts were obtained under non-denaturing conditions. A series of HR-SPROX analyses were carried out with GdmCl concentrations ranging from 0 to 3 M (Figure 3A). The experiments included five technical replicates obtained from two biological replicates, with each technical replicate being exposed to 30 denaturant concentrations and tagged with three different sets of 10-plex TMTs prior to LC-MS/MS analysis. Together, the data provided quantitative information for 10,412 unique methionine-containing peptides (in unoxidized and/or oxidized forms) mapped to 3,158 protein groups (Table 1, Supplementary Tables 1-3). All raw and processed data have been deposited to the ProteomeXchange Consortium via the PRIDE (32) partner repository with the dataset identifier PXD011456.

**Figure 3.**
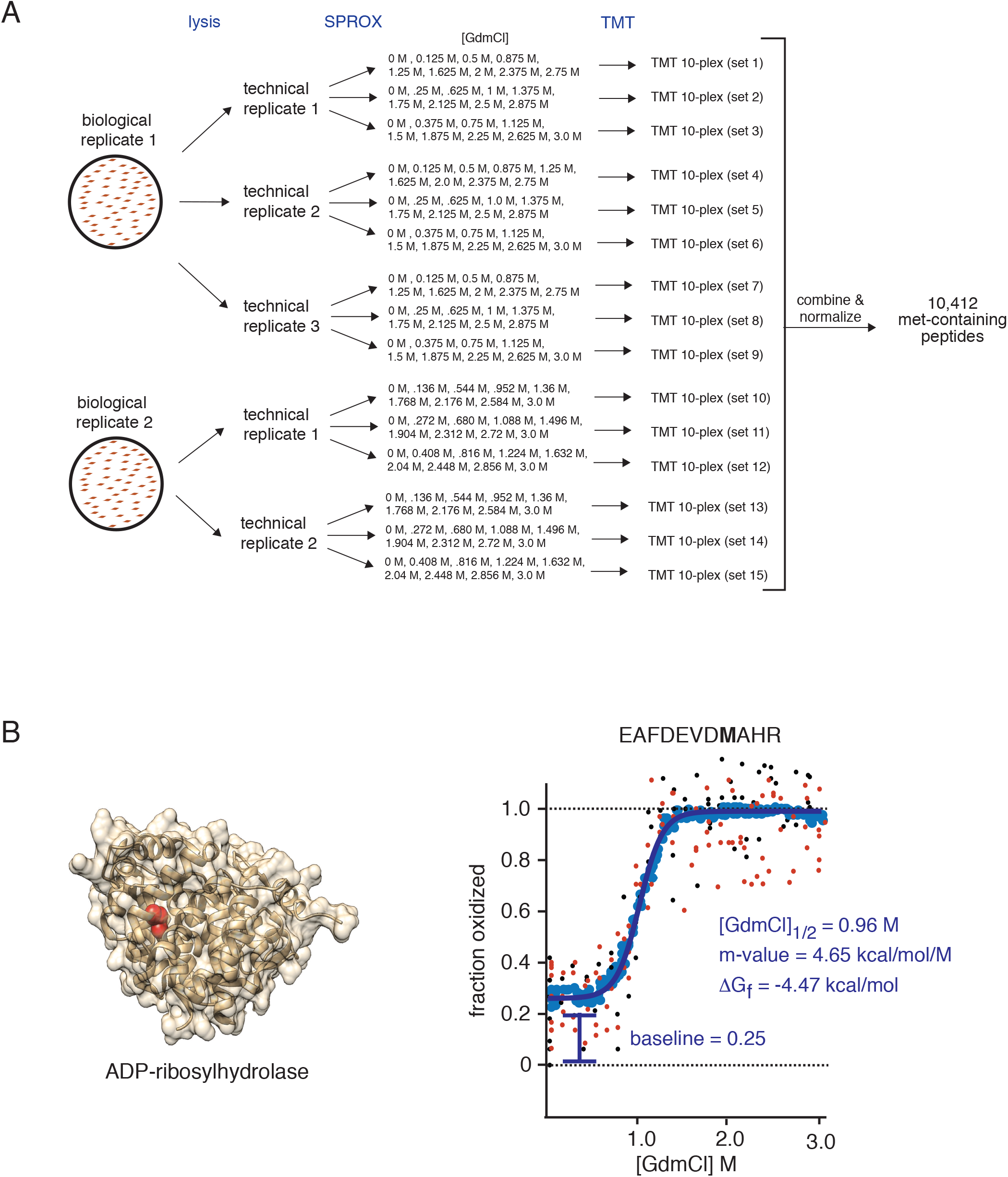
Experimental design and example data for proteome-wide HR-SPROX analysis. (A) Schematic representation of the experimental design for HR-SPROX analysis of the human fibroblast proteome indicating the replicates and denaturant concentrations used in the study. Each technical replicate consists of 3 complementary sets of GdmCl concentrations tagged with TMT 10-plex reagents. The data were combined after normalization, providing denaturation curves for 10,412 met-containing peptides. (B) Example HR-SPROX data for a protected methionine (highlighted in red) within a tryptic peptide in ADP-ribosylhydrolase (2FOZ). By combining a number of replicate measurements, detailed denaturation curves were obtained. Small black and red points indicate data obtained from methionine and methionine sulfoxide-containing peptides, respectively. The plots were smoothed by obtaining moving averages of the data (blue points) and fit by a two-state model (blue line) to determine ΔG_folding_, m-values, [GdmCl]_1/2_ and baseline oxidation values. Also see Figures S1–3.

**Table 1.** Combined quantified proteome coverage of HR-SPROX analyses (-TMAO)

| **incorporated residue** | **unique peptides** | **unique proteins** |
| --- | --- | --- |
| met(unmodified) | 5,719 | 1,977 |
| met(ox) | 9,350 | 2,977 |
| met(unmodified) & met(ox) | 4,565 | 1,745 |
| met(unmodified) or met(ox) | 10,412 | 3,158 |

The H_2_O_2_ concentrations and oxidation times used in our experiments were sufficient to oxidize exposed methionines to methionine sulfoxides (Met-O) but did not lead to the formation of significant levels of methionine sulfones (Met-O_2_) (Figure S1). The only other oxidized residue that was detectable at appreciable levels was sulfonic acid (Cys-O_3_). However, levels of sulfonic acid-containing peptides were significantly lower than methionine sulfoxide-containing peptides and their formation did not result in any calculation bias in measurements of methionine oxidation (Figure S1).

We initially analyzed methionine oxidation levels in individual experiments and showed that there was a significant correlation between replicate experiments (Figure S2). Additionally, there was a significant correlation between denaturation curves of individual peptides independently obtained from disappearance of methionines and appearance of methionine sulfoxides (Figure S2). Given that replicate experiments were in general agreement with each other, we combined all available data (all replicates, encompassing measurements of oxidized and/or unoxidized peptides based on availability) to obtain high resolution denaturation curves for individual peptides. After merging datasets, the number of denaturant concentration points analyzed for individual methionine-containing peptides ranged from 10 to 300, with a median of 50 (Figure S3).

As an example, Figure 3B illustrates a HR-SPROX denaturation curve for a protected methionine residue in ADP-ribosylhydrolase. We noticed that although individual TMT data points for a given peptide generally overlapped and had consistent trends, outliers were typically present and heavily biased subsequent least-squares regression analyses of the denaturation curves. We therefore obtained moving averages of the denaturation data and used the resulting smoothed plots to conduct least squares regression analyses (Figure 3B). The baseline oxidation, [GdmCl]_1/2_, m-value and ΔG_folding_ measurements for individual methionine-containing peptides are tabulated in Supplementary Table 4.

### Baseline methionine oxidation levels correlate with secondary structure and solvent accessibilities

We first focused on analysis of baseline oxidation levels in the absence of denaturant. The data indicated that the distribution of baseline oxidation levels within the proteome is remarkably bimodal, with approximately half of the methionines forming a relatively well-protected population and half forming a poorly protected population (Figure 4A). Based on these measurements, we set out to identify sequence and structural parameters that correlate with methionines’ susceptibility to oxidation.

**Figure 4.**
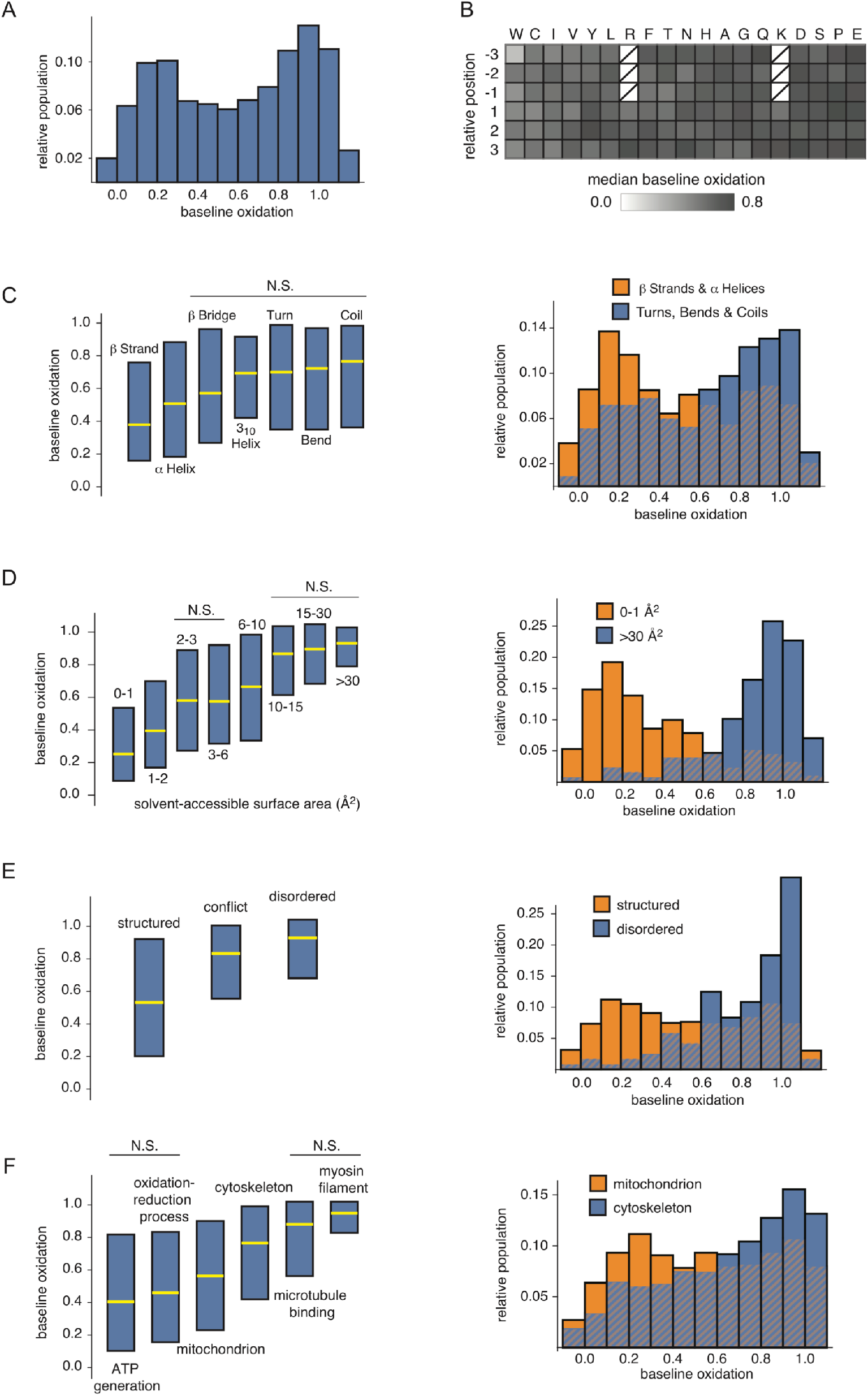
Global analysis of methionine baseline oxidation. (A) The distribution of fractional methionine baseline oxidation levels within the analyzed proteome. Note that due to experimental noise, it is possible for +H_2_O_2_/0M GdmCl reporter ions to have a higher intensity than +H_2_O_2_/3M GdmCl reporter ions, resulting in measured oxidation levels greater than 1. (B) Relative baseline oxidation levels of peptides with different methionine-neighboring residues. Note that because the data consist of tryptic peptides, sequences having lysine and arginine residues on the N-side of methionines are rare and not included in the analysis. (C-F) Comparison of baseline oxidation levels of methionines contained within different DSSP secondary structure elements (C), having different solvent accessible surface areas (D), having different disordered designations in accordance with MobiDB (E), and contained within proteins mapped to different gene ontologies (F). For all box plots, the medians are represented by yellow lines, and blue boxes denote interquartile ranges. The distributions being compared in box plots significantly differ from one another with p-values less than 0.001 using the Mann Whitney U test, with the exception of those grouped together under bars labeled “N.S.” The histograms indicate the distributions of selected categories. Cross hatching indicates overlapping bars. Also see Figures S4–5.

We did not find strong correlations between methionine oxidation and identities of neighboring residues in the primary sequence, beyond slightly decreased oxidation levels near hydrophobic residues (Figure 4B, S4). As discussed below, we can rationalize this trend by considering that hydrophobic residues are more likely to be in the protein core where methionines would lack exposure to solvent and thus be less susceptible to oxidation. Alternatively, as has been noted previously, the direct interaction of the methionine sidechain with aromatic residues may decrease the reactivity of the sulfur atom with oxygen (33).

We next cross-referenced baseline oxidation levels to the secondary structure context of the methionine residues using the Define Secondary Structure of Proteins (DSSP) analysis of corresponding protein databank (PDB) structures (34). Structural data mapped to 1,016 proteins in our dataset indicated that highly oxidizable methionines tend to be contained in turns, bends and unstructured coils, whereas more protected methionines tended to be in rigid secondary structure elements such as β-strands and a-helices (Figure 4C, Supplementary Table 5). We subsequently used PDB structures to measure the solvent accessible surface area (SASA) of methionines using the VMD molecular visualization program (35). We observed a very strong positive correlation between the SASA of a methionine residue and its susceptibility to oxidation (Figure 4D, Supplementary Table 5). Together, the data provide proteome-wide evidence that secondary structure context and solvent accessibilities of methionine residues are the primary determinants of their susceptibility to oxidation. This trend had been previously noted for a limited number of individual proteins (27).

### Intrinsically disordered proteins are highly prone to oxidation in cell extracts

A large number of protein regions and domains have been classified as *intrinsically disordered,* primarily based on lack of evidence for structure in crystallography or NMR studies, or low complexity patterns of amino acid usage in their primary sequence (36). However, it is unclear what fraction of these sites retain their disorder in the complex intracellular environment of the cell where they could potentially be interacting with binding partners and other stabilizing factors. Given the strong correlation between localized structure and methionine oxidation established above, we assessed the baseline oxidation levels of methionines within regions classified as *disordered* in a curated database of intrinsically disordered proteins (MobiDB) (37). Our analysis indicated that methionines within regions classified as *disordered* are significantly more prone to oxidation than those classified as *structured* (Figure 4E, Supplementary Table 5). This suggests that as a group, proteins that have been classified as *intrinsically disordered* mostly retain this property in crude extracts where they could potentially be interacting with stabilizing factors and binding partners. Thus, methionine oxidation rates can potentially help resolve ambiguities where conflicting evidence exists regarding the *structured/disordered* designation of a protein region.

### Susceptibility to oxidation is enriched in specific gene ontologies

We conducted gene ontology (GO) enrichment analyses to identify GO categories that were statistically enriched in protected or exposed methionines (Figure 4F, S5, Supplementary Table 6). GO categories most enriched in protected methionines included mitochondrial proteins, as well as proteins involved in oxidation-reduction reactions and ATP synthesis. It is notable that these categories contain proteins that are particularly exposed to reactive oxygen species (ROS) within the cell. It is possible that the prevalence of protected methionines in these proteins is an adaptive response to safeguard against ROS-induced oxidative damage. Among the GO categories most enriched in exposed methionines were cytoskeletal proteins. It has been shown that enzyme-mediated methionine oxidation is a regulatory mechanism for cytoskeletal dynamics. MICAL proteins are F-actin disassembling factors that modify the actin cytoskeleton by oxidizing two conserved methionine residues in actin (38, 39). Our data indicate that not only are MICAL-targeted methionines highly prone to oxidation (Figure 5A), but that as a group, cytoskeletal proteins contain an atypically large number of exposed methionines that could potentially be targetable by other oxygenases (40).

**Figure 5.**
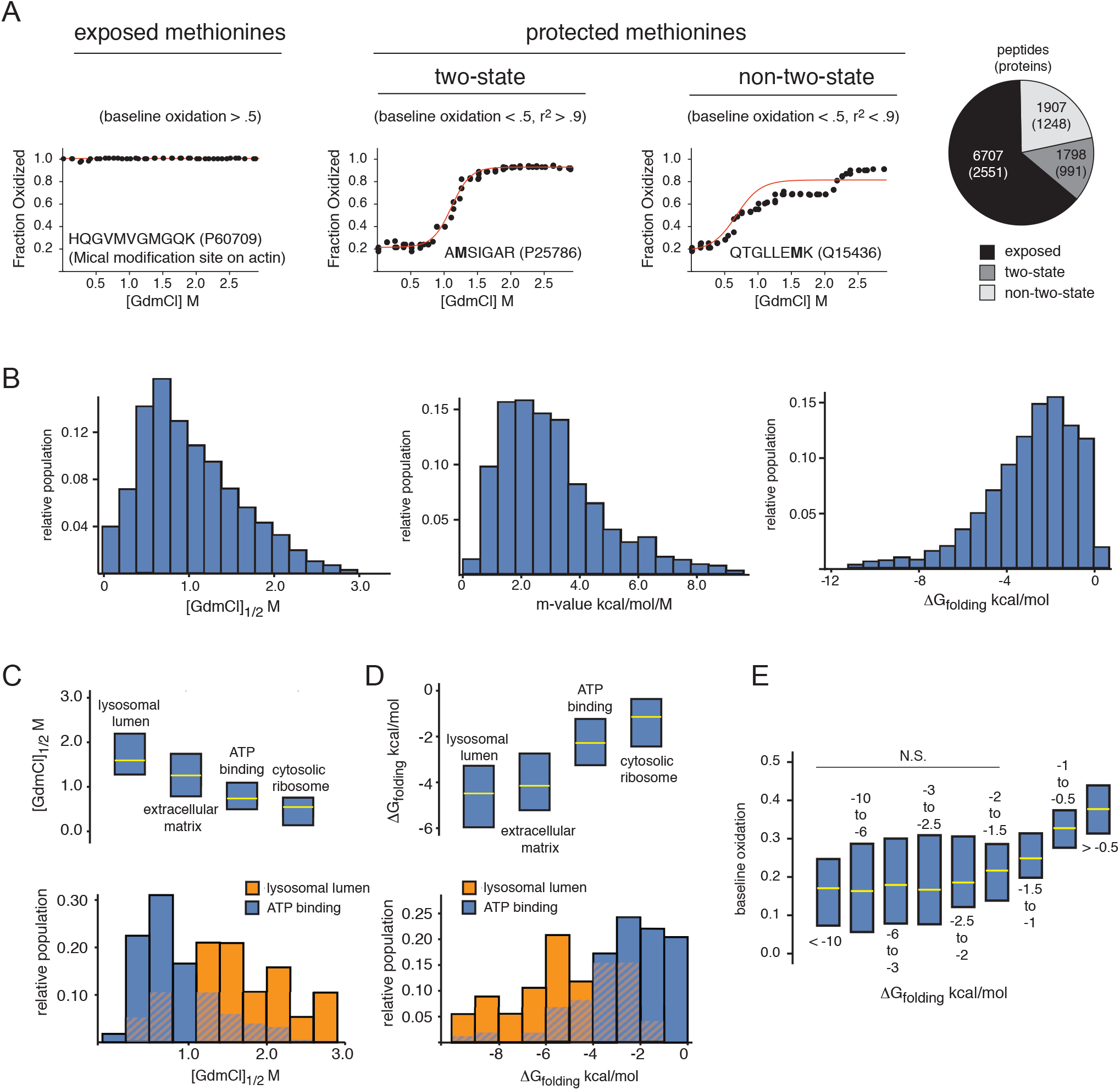
Global analysis of thermodynamic folding parameters. (A) Quantified methionine-containing peptides were separated into one of three categories based on the observed characteristics of their denaturation curves: solvent exposed methionines (baseline oxidation>.5), protected two-state proteins (baseline oxidation<.5 and goodness-of-fit r^2^ to two-state model>.9) and protected non-two-state proteins (baseline oxidation<.5 and r^2^<.9). The scatter plots show averaged denaturation curves for example peptides in each category. The peptide used as an example of an exposed methionine is the MICAL oxidation site on actin, discussed in text. The pie chart indicates the number of peptides and proteins in each category. (B) Distributions of [GdmCl]_1/2_, m-values and ΔG_folding_ measurements for two-state peptides. (C-D) Distributions of [GdmCl]_1/2_ and ΔG_folding_ measurements for example gene ontology terms enriched in stable or unstable methionine-containing peptides. (E) The relationship between protein stability (ΔG_folding_) and baseline methionine oxidation levels. See Figure 4 for description of box and bar plots. Also see Figure S6.

### HR-SPROX uncovers proteome-wide distributions of [GdmCl]_1/2_, ΔG_folding_ and m-values

We next focused on analyzing thermodynamic folding parameters based on peptide HR-SPROX denaturation curves. We limited our subsequent analyses to peptides that contained protected methionines (conservatively defined as having a baseline oxidation levels less than 0.5) and were well fit by a two-state folding model (r^2^>0.9). This “two-state” subset consisted of 1,798 peptides mapped to 991 proteins (mean peptide to protein ratio of 1.7) (Figure 5A). Peptides that contained protected methionines but were poorly fit by a two-state folding model (r^2^<0.9) were designated as “non-two-state”. These peptides fell into two categories. The first were peptides that showed poor fit to the two-state model due to low signal to noise (e.g. low intensity peptides). The second were peptides that displayed multi-phasic unfolding patterns, perhaps due to the presence of partially folded intermediates populated during the course of denaturation. However, the resolution and signal to noise level of our data were not sufficient to accurately quantify the thermodynamic properties of folding intermediates in the latter category of peptides.

For “two-state” peptides, the distributions of ΔG_folding_, m-values, [GdmCl]_1/2_ measurements are plotted in Figure 5B. The median values for these three parameters were −2.6 kcal/mol, 2.8 kcal/mol/M and 0.91 M respectively. As a whole, our measured stabilities and m-values appear lower than the majority of protein folding models studied to date and theoretical proteome-wide predictions for global protein stabilities (29, 41). It is therefore likely that for most proteins, HR-SPROX measurements do not reflect the global stability of the protein, but rather sub-global stabilities of localized cooperative unfolding units encompassing the methionines. In this way, the exposure of methionines to oxidation through sub-global unfolding may be similar in nature to native hydrogen deuterium exchange (HDX) experiments (42).

### Protein stability is elevated in specific gene ontologies

We conducted gene ontology (GO) analyses to identify protein categories that are statistically enriched in highly stable or highly unstable regions as determined by [GdmCl]_1/2_ and ΔG_folding_ measurements of their methionine residues (Figure 5C-D, S6, Supplementary Table 6). Among the most stable subsets of the proteome were proteins localized to the lysosomal lumen and extracellular space. Unfolding of target proteins in the denaturing environment of the lysosome is known to be required for their hydrolysis (43). It is therefore likely that resident proteins of the lysosome have evolved to be resistant to denaturation. Similarly, proteins localized to the oxidizing environment of the extracellular space are enriched in stabilizing disulfide bonds.

Among the least stable subsets of the proteome were ATP binding proteins and ribosomal proteins. At ATP concentrations present in cell extracts, many ATP binding proteins may exist in ATP-unbound forms that are predicted to be less stable than the ATP-bound forms (44). At higher ATP concentrations, the binding equilibrium may shift to the bound form, resulting in ligand-induced stabilization. Similarly, ribosomal proteins are stabilized by stoichiometric association to form the intact small and large subunits of the ribosome. During the course of chemical denaturation, as the higher order structure of the ribosome is disrupted, orphaned ribosomal proteins appear to be relatively thermodynamically unstable.

### Protein stability impacts the rate of oxidation for a subset of the proteome

We next analyzed the relationship between folding stability and baseline methionine oxidation (Figure 5E). The data indicated that ΔG_folding_ is not a strong predictor of baseline susceptibility to oxidation for two-state proteins that are relatively stable (ΔG_folding_ < −1.5 kcal/mol). However, once protein stability declines beyond a minimum threshold, baseline methionine oxidation rates begin to rise. Thus, in unstable protein regions, oxidation from the unfolded state is the dominant pathway for oxidation, and the rate of oxidation is contingent on the folding equilibrium. Conversely, in stable protein regions, where the folding equilibrium is heavily shifted towards the native state, oxidation occurs primarily from the folded structure. In this regime, the rate of oxidation is independent of the folding equilibrium and instead appears to be limited by the solvent accessibility of the methionine in the native state.

### HR-SPROX enables analysis of inter-and intra-domain variability in stability

An important feature of the HR-SPROX analysis is that it can potentially provide localized stability information within proteins. In theory, different methionines within the same protein provide independent and localized stability probes. In some multi-domain proteins, each domain may act as an independent cooperative unfolding unit (45). For these proteins, different methionines within the same domain are expected to report similar stabilities. To assess this feature of our data, we examined intra-domain variability in our stability measurements. In general, we found that different methionines within the same domain have similar stabilities (Figure 6).

**Figure 6.**
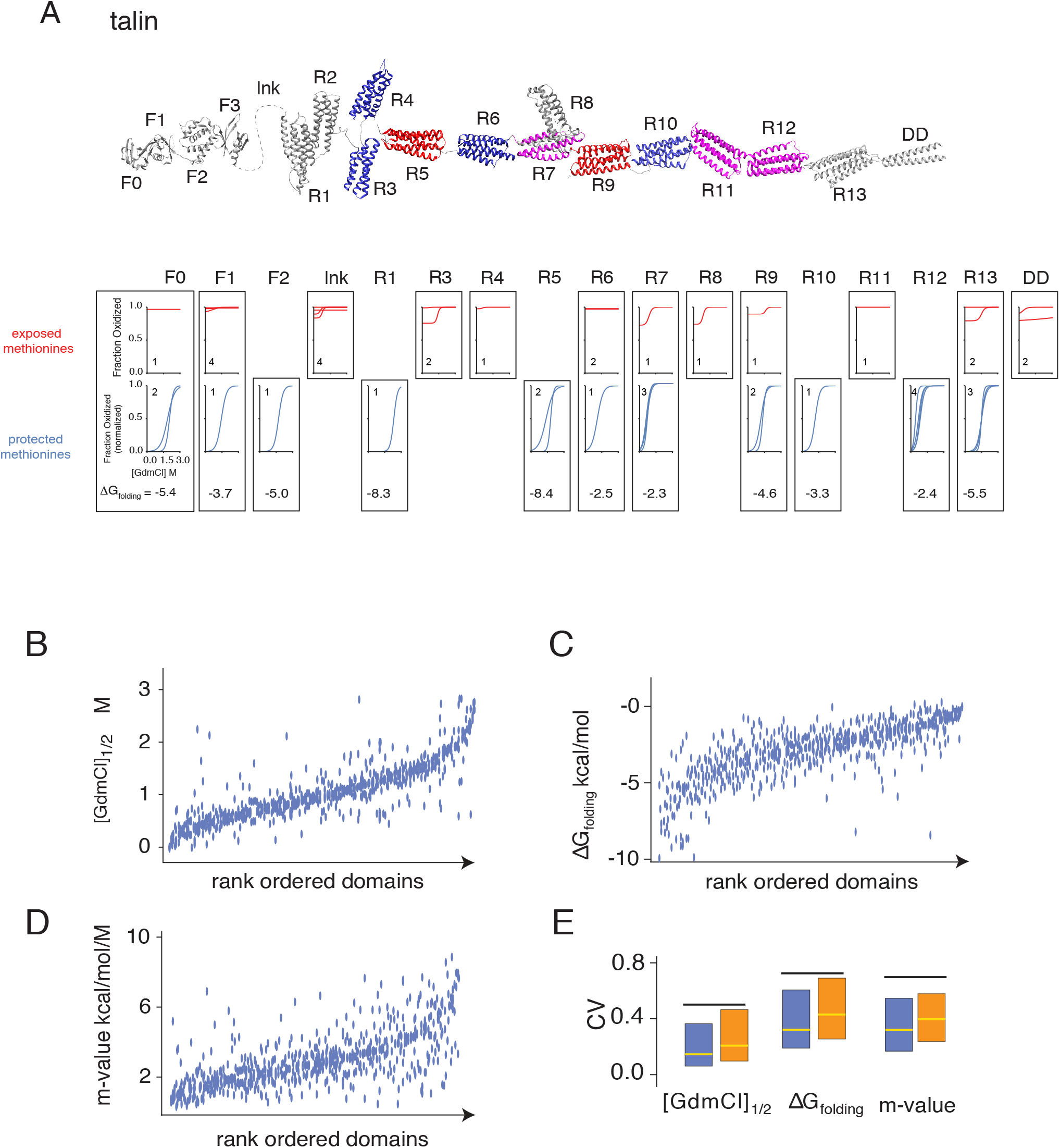
Inter- and intra-domain comparisons of thermodynamic folding parameters. (A) Talin is presented as an example of an analyzed multi-domain protein. The depicted structure is a composite of available structures for individual domains. The color coding indicates “weak” (blue), “intermediate” (pink) and “strong” (red) mechanical stabilities designated by Haining et al.(46) based on unfolding force magnitudes of domains in smAFM and steered molecular dynamics studies. Denaturation curves represent exposed (red) or protected (blue) methionine-containing peptides in different domains. For protected methionines, fractional oxidation measurements were normalized with respect to the native and denatured baselines. The reported ΔG_folding_ is the median of all protected methionines. The inset numbers indicate the number of peptide denaturation curves contained within plots for each domain. (B-D) Global analysis of intra-domain variation. Using boundaries defined by the Pfam database, [GdmCl]_1/2_ (B), ΔG_folding_ (C), and m-values (D) were assigned to specific domains. Measurements within 221 domains that contained more than one methionine were rank ordered based on their median values. Peptide measurements mapped to a given domain are plotted on the y-axis for each rank-ordered domain represented on the x-axis. (E) The distributions of coefficients of variation (CV) of [GdmCl]_1/2_, ΔG_folding_ and m-value measurements for different peptides within individual domains (intra-domain variation, blue box plots), within individual proteins (intra-protein variation, orange box plots), and CV of the measurements in the entire dataset (intra-proteome variation, black lines). See Figure 4 for description of box plots. The blue dots represent the median CV values for all analyzed peptides in the proteome.

An example of this trend is shown for the cytoskeletal protein talin (Figure 6A). Talin is a large, amphipathic multi-domain protein that forms a mechanical linkage between integrins and the cellular cytoskeleton. The interaction between talin domains and vinculin in focal adhesions is force-dependent and is thought to occur through mechanical unfolding of specific talin domains that expose cryptic vinculin binding sites. To identify potential binding sites, mechanical stabilities of individual talin domains have been previously investigated by single molecule atomic force microscopy (smAFM) (46-48). Here, we were able to track the stability and solvent exposure of 16 of the 18 domains in talin. Our data are largely consistent with the smAFM findings. We found that domains shown to have low mechanical stability by smAFM, including the linker region, R3 and R4 are solely composed of highly exposed methionines. Conversely, domains that have been shown to be highly stable, such as R5 and R9 contained protected and stable methionines. Additionally, our analysis provided thermodynamic data on domains that have not been previously studied by smAFM. For example, we identified R13 as another highly stable talin domain. These results highlight the ability of HR-SPROX to analyze localized thermodynamic stability data for large multi-domain proteins that are difficult to study by traditional protein denaturation experiments.

Within individual talin domains where multiple methionine-containing peptides could be identified, the stability trends were in good agreement with each other (see overlap between protected methionine curves in Figure 6A). To examine the generality of this trend, we used the Pfam database of protein domains (49) to globally define domain boundaries within our dataset and analyzed intra-domain variability in our measurements. The analysis indicated that variabilities in [GdmCl]_1/2_, ΔG_folding_ and m-values were significantly lower within domains in comparison to variabilities of the parameters within the overall proteome (Figure 6B-E). The results are consistent with the idea that many domains behave as independent cooperative unfolding units.

### The chemical chaperone TMAO has variable stabilizing effects on the proteome

To determine whether HR-SPROX is capable of monitoring changes in protein stability induced by environmental alterations, we repeated our analyses in the presence of the chemical chaperone trimethylamine N-oxide (TMAO). TMAO is an amine-oxide osmolyte (50). It is found in the tissues of a variety of marine organisms where it is thought to counteract the destabilizing effects of urea and pressure on proteins. It has been postulated that TMAO stabilizes proteins by increasing the free energy of the unfolded state through solvent exclusion effects, thus shifting the folding equilibrium towards the native state (51, 52). Although the stabilizing effects of TMAO had previously been observed for a number of proteins (53, 54), the selectivity of its stabilizing effects have not been investigated on a proteome-wide scale. We conducted a limited HR-SPROX analysis in the presence of 1 M TMAO (Figure S7) and compared the baseline oxidation levels, [GdmCl]_1/2_, ΔG_folding_ and m-values of methionine-containing peptides to untreated controls (Figure 7, S7). Overall, we observed that TMAO decreased baseline oxidation levels and increased the stability of the proteome, although the magnitudes of these effects were variable between proteins (Figure 7A).

**Figure 7.**
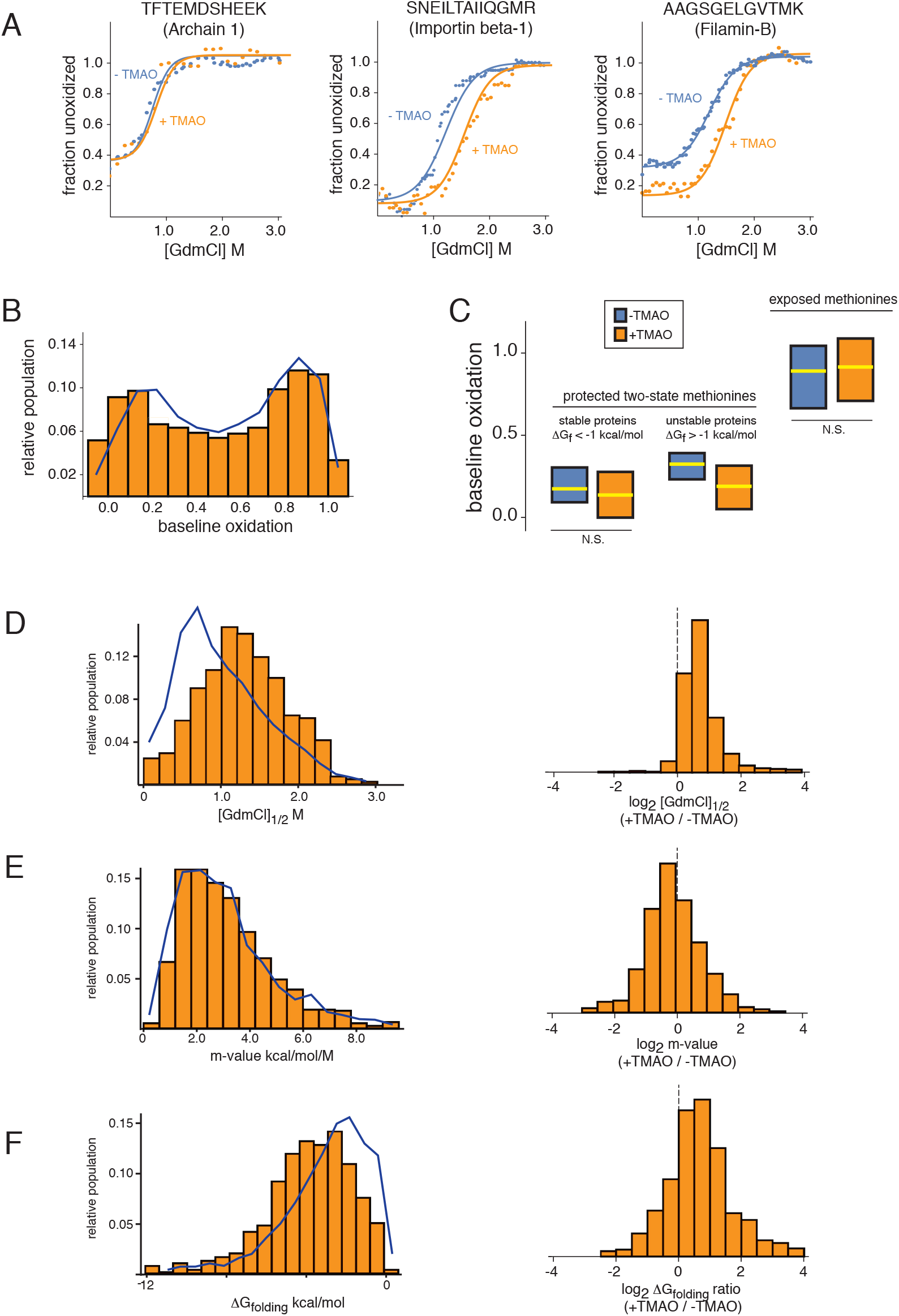
The effect of the chemical chaperone TMAO on proteome stability. (A) The effects of TMAO on denaturation curves of three example peptides mapped to different proteins. The data indicate that TMAO has variable effects on baseline oxidation levels and stabilities of different protein regions. (B) The distribution of relative methionine baseline oxidation levels within the analyzed proteome in the absence (blue lines, reproduced from Figure 4) and presence of 1 M TMAO (orange bars). (C) Impact of TMAO on baseline oxidation levels of exposed and protected proteins with different stabilities. (D-F) Impact of TMAO on distributions of [GdmCl]_1/2_ (D), m-values (E) and ΔG_folding_ (F). The distributions of the measurements are compared in the absence (blue lines, reproduced from Figure 5) and presence of 1 M TMAO (orange bars), and also presented as log2 ratios of the measurements. See Figure 4 for description of box plots. Also see Figure S7.

Our data indicated that TMAO decreased baseline oxidation levels of protected methionines within unstable proteins (ΔG_folding_>−1.0 kcal/mol), but that this effect was diminished for more stable proteins (Figure 7B-C). This trend is consistent with the idea that unlike stable protein regions, methionine oxidation in unstable protein regions occurs primarily from the unfolded state. Hence, modulation of the folding equilibrium by a chemical chaperone can effectively decrease oxidation rates in unstable protein regions (see above and Figure 5E). However, TMAO did not significantly decrease the oxidation rates of most highly exposed methionines, indicating that most unstructured regions within proteins (including regions with non-rigid secondary structures, high solvent accessibility and those characterized as *intrinsically disordered*) do not transform to stably folded structures in the presence of 1 M TMAO (Figure 7C, S7). Although our experiments did not investigate the efficacy of TMAO at higher concentrations, the results indicate that at levels where it has a significant impact on the stability of structured proteins (Figure 7D-F), TMAO is not able to impose structure on the majority of disordered regions within the proteome.

## DISCUSSION

In this study, we investigated the thermodynamic folding stability of the human proteome using methionine oxidation as a probe. Our data indicate that rates of methionine oxidation are correlated to localized structure and amenable to detection and quantitation by bottom-up proteomics. However, this approach has at least two important limitations. First, the analysis is limited to tryptic peptides that contain methionines, which is one of the least abundant residues in natural proteins. For example, in our analyses, although we were able to detect a total of 60,776 unique peptides in our combined experiments, only 19% contained methionines and could be analyzed by HR-SPROX. Second, methionine oxidation is not a benign modification and may in itself impact folding stability. Thus, oxidation of one methionine could artifactually alter the localized stability of a second methionine located elsewhere in the same protein. This effect could potentially skew ΔG_folding_ and m-value measurements and mask multi-phasic unfolding behavior. Nonetheless, methionine oxidation likely represents a significantly less disruptive modification in comparison to alternative proteomic approaches for analysis of protein structure based on limited proteolysis or indiscriminate radical footprinting.

Using HR-SPROX we were able to quantify the susceptibility of a large number of methionines to oxidation by H_2_O_2_. Our results indicate that the distribution of methionine oxidation susceptibilities is strikingly bimodal. Methionines within the human proteome appear to be divided into two roughly equal-sized groups: solvent exposed methionines that are highly prone to oxidation, and buried methionines that are protected from oxidation. Methionines are hydrophobic residues that would be expected to reside primarily in the cores of proteins based solely on their chemical properties. However, it has been suggested that exposed surface methionines can play a protective role by acting as sinks for reactive oxygen species (ROS) (55). Within the cell, the oxidation of methionines is enzymatically reversible and their reduction by methionine sulfoxide reductases (MSRs), and subsequent formation of NADPH, forms a complete redox cycle (56, 57). Thus, reversible oxidation of surface methionines can act as a removal mechanism for cellular ROS and prevent the irreversible oxidation of other biomolecules (55). In support of this idea, our data provides global experimental evidence for the high prevalence of exposed oxidation-prone methionines in the human proteome.

Our analysis was able to measure thermodynamic folding stabilities (ΔG_folding_) for ~1,800 protein regions that contain highly protected methionines and unfold in accordance with a two-state model. To the best of our knowledge, this analysis represents the largest survey of thermodynamic protein stabilities conducted to date. The data highlight the remarkable range of stabilities within the proteome and suggest that stability is a physical property that has adapted to meet the specific functional needs of proteins. For example, our data highlight lysosomal and extracellular proteins as two subsets of the proteome that have evolved high stabilities, likely due to the physical demands of their local environments.

Our proteome-wide data also uncovered an important relationship between folding stability and methionine oxidation. We showed that in general there are two oxidation pathways for buried methionine residues. Oxidation can occur rapidly from the denatured state or more slowly from the folded state. Most buried and protected methionines reside in structured regions with stabilities that are high enough to heavily favor oxidation through the native state. However, ~10% of protected methionines analyzed in our study were located in regions that were sufficiently unstable (ΔG_folding_>−1.5 kcal/mol) to favor oxidation through the denatured state. The oxidation rate of this subset of the proteome is contingent on folding stability and can be diminished by stabilizing factors such as chemical chaperones (e.g. TMAO).

It has long been known that TMAO stabilizes proteins in saltwater fish, sharks, crustaceans and molluscs (58). It is thought that this activity may counteract the destabilizing effects of urea and pressure in deep-sea animals. However, the exact physical mechanism of the stabilizing effects of TMAO remains under active investigation. It has been shown that interactions between TMAO and the peptide backbone raises the energy of unfolded structures and energetically favors the folded state (51, 59). Here, we have shown that TMAO does indeed stabilize much of the proteome. However, the magnitude of its effect is variable between proteins and domains. Our experiments have quantified the effects of TMAO on diverse protein substrates and the results may provide a useful resource for evaluating putative mechanisms of stabilization by this osmolyte. More generally, this study demonstrates that HR-SPROX can be used to identify client-sets of chaperones by globally quantifying their effects on protein folding stabilities.

It has been predicted that ~30% of the human proteome contains regions that can be classified as intrinsically disordered (IDPs) (11). However, investigating the prevalence and properties of these regions within the cellular environment has proven challenging. In theory, the presence of interacting partners and molecular chaperones in cell extracts may impose structure on regions that otherwise appear disordered in purified systems. Here, we used HR-SPROX to probe the structure and stability of regions classified as IDPs in crude extracts. Our data indicate that in comparison to the rest of the proteome, methionines in IDPs are highly prone to oxidation. These results suggest that most proteins designated as IDPs remain relatively unstructured even in crude extracts.

Our data also indicate that the addition of TMAO does not alter the oxidation properties of most IDPs, suggesting that most disordered regions can not be significantly folded by the addition of 1 M concentrations of this osmolyte. As described above, TMAO is thought to modulate the equilibrium constant between the unfolded and the native state of stably folded proteins. However, this mechanism of action is contingent on the existence of localized energy minima representative of denatured and native states. Our data suggest that the folding energy landscapes of most IDPs may not contain localized minima representative of unique native states. Thus, IDPs may represent fundamentally different structures in comparison to “unstable proteins”. While the folding equilibrium of the latter can be altered in favor of a native state by the addition of osmolytes such as TMAO, the former inherently lack a native structure and are incapable of achieving a singular folded state even in the presence of stabilizing agents.

Together, our data validate a global methodology for quantitative analysis of protein stability and provide a proteome-wide census of ΔG_folding_ for the human proteome. This approach may prove useful in investigating proteome-wide trends in folding stabilities and can be employed in future studies to investigate the stabilizing effects of molecular chaperones, characterize the impact of environmental stresses on protein folding, and examine cross-species conservation of folding thermodynamics.

## SUPPLEMENTAL INFORMATION

Supplemental information include detailed experimental methods, seven figures and seven data tables.

## ACKNOWLEDGEMENTS

We thank Dr. Michael Fitzgerald, Dr. Mark Dumont and the members of the Ghaemmaghami and Fu labs for helpful comments on the manuscript. This work was supported by grants from the National Science Foundation (MCB-1350165 CAREER) and National Institutes of Health (R35 GM119502-01, T32 GM068411, 1S100D021486-1, 1S10OD025242-01).

## SUPPLEMENTARY INFORMATION

### METHODS

#### Cell culture

Human diploid fibroblasts expressing telomerase (HCA2-hTert) (31) were cultured at 37°C, 5% CO_2_, in Eagle’s minimum essential medium (EMEM) supplemented with 15% fetal bovine serum (FBS), 100 U/mL penicillin and 100 U/mL streptomycin. Cells were grown to 100% confluency and maintained for at least 4 days to ensure they reached a quiescent state. Cells were harvested, washed twice with phosphate buffered saline (PBS), and the pellets were stored at −80^°^C until analysis.

#### Preparation of stock solutions

Native lysis buffer was prepared by dissolving 1 Pierce EDTA-free protease inhibitor mini tablet in 10 mL of a filtered 20 mM sodium phosphate buffer (pH 7.4) with 50 mM NaCl (Fisher). All GdmCl stocks were prepared volumetrically by mixing ratios of a filtered 7 M stock solution of ultrapure GdmCl (Sigma-Aldrich) dissolved in 20 mM sodium phosphate buffer (pH 7.4) with filtered phosphate buffer only. All GdmCl stocks were kept at room temperature (RT) for no longer than 3 months. Urea-containing buffers were prepared fresh on the day of analysis and made volumetrically from high purity (≥99.5%) urea pellets (Sigma-Aldrich) for a final concentration of 8 M urea in 0.1 M triethylammonium bicarbonate (TEAB) buffer (pH 8.5). tris(2-carboxyethyl)phosphine (TCEP) quenching buffers were prepared volumetrically from TCEP hydrochloride (Sigma-Aldrich), adjusted to neutral pH with concentrated NaOH (Fisher), and diluted to 0.5 M (of note, for samples 1-10 the pH was adjusted only by resuspending TCEP at 0.5 M in TEAB buffer). Iodoacetamide (IAA) buffer for protein alkylation was prepared immediately before use from high purity IAA (Sigma) dissolved in urea buffer. For all buffers, MS-grade water (Thermo Scientific) was used as a diluent.

#### Protein extraction

On the day of analysis, cell pellets were resuspended in native lysis buffer at 50 μL per 10^6^ cells. Lysis was accomplished by repeated passage through a 16-gauge sterile syringe, resting on ice, after which the samples were centrifuged for 10 min at 13000xg while at 4°C, and the supernatants collected. Protein concentration was subsequently measured using a BCA assay and samples were diluted to a final concentration of 5 mg/mL, with the exception of experiments 10-15 which were diluted to 2 mg/mL.

#### Sample preparation and TMT tagging

Protein extracts were aliquoted out in 25 μL portions into 30 wells of a 96-deepwell polypropylene microplate (Fisher Scientific). 3 wells were kept as a control (no H_2_O_2_, no GdmCl) for measuring background levels of methionine oxidation, and the remaining wells were split into 3 complementary sets of denaturant-containing buffers such that their final concentrations ranged between 0-3 M GdmCl (see Figure 3A), allowed to equilibrate for 1 hr at RT, and oxidized with 3% H_2_O_2_ for 1 min. Oxidation was quenched using a four-fold molar excess of TCEP, and protein was purified using 30-kDa centrifugal filters (Millipore Sigma) in a modified iFASP procedure (60). During this process, proteins were fully denatured using 8 M urea, disulfide bonds were reduced with 5 mM TCEP in urea buffer for 1 hr at RT, and proteins were alkylated by incubating with 50 mM IAA for 20 min at RT in darkness. After washing out the denaturant, proteins were digested with trypsin protease (Pierce) at a 1:100 ratio (g protein/g trypsin), incubating overnight at 37°C in a wet bath. Trypsin was quenched by adding formic acid (Fisher Scientific) to a final concentration of 1%, after which samples were dried down and 25 μg (10 μg for experiments 10-15) of digest was resuspended in 0.1 M TEAB for labeling using TMT 10-plex reagents (Thermo Scientific). TMT labeling was accomplished using 0.2 mg of TMT 10plex reagents (Thermo Scientific) suspended in acetonitrile (Thermo Scientific) and incubating for 1 hr at RT. TMT labeling was quenched using 5% hydroxylamine (Fisher Scientific), incubating for 15 min at RT. For each of the TMT sets, 5 μg portions of labeled digest were combined and dried down.

#### Peptide fractionation by spin column

To increase coverage, samples were fractionated using lab-made C18 spin columns. Columns were first conditioned using acetonitrile (ACN) and 100 mM ammonium formate (AF) buffer (pH 10). Samples were resuspended in 50 μL of AF buffer and added to the columns. After washing the columns with 5% ACN in AF buffer, samples were eluted with 10%, 12.5%, 15%, 17.5%, 20%, 22.5%, 25%, and then 50% ACN (in AF buffer). Eluent fractions were then dried down and resuspended in MS-grade 0.1% trifluoroacetic acid (TFA) in water (Thermo Scientific) at a concentration of 0.25 mg/ml for LC-MS/MS analysis.

#### Proteome stability in the presence of TMAO

For these analyses, protein extracts were prepared at a concentration of ~5 mg/ml, using the method described above. On the day of analysis, a concentrated TMAO stock solution was prepared from TMAO dihydrate dissolved in 20 mM sodium phosphate buffer (pH 7.4). Immediately prior to incubation in GdmCl, samples were mixed with the TMAO stock solution such that the final concentration of TMAO was 1 M. Samples were then processed and fractionated as above.

#### Mass spectrometric analysis

Analyses of -TMAO experiments 1-6 and +TMAO experiments 1-3 (see Figure 3A, S7A) were conducted on a Q Exactive Plus instrument, and the remainder of the experiments were conducted on a Fusion Lumos Tribrid instrument as described below.

For Fusion Lumos LC-MS/MS analysis, peptides were injected onto a homemade 30 cm C18 column with 1.8 um beads (Sepax), with an Easy nLC-1200 HPLC (Thermo Fisher), connected to a Fusion Lumos Tribrid mass spectrometer (Thermo Fisher). Solvent A was 0.1% formic acid in water, while solvent B was 0.1% formic acid in 80% ACN. Ions were introduced to the mass spectrometer using a Nanospray Flex source operating at 2 kV. The gradient began at 3% B and held for 2 min, increased to 10% B over 7 min, increased to 38% B over 94 min, then ramped up to 90% B in 5 min and was held for 3 min, before returning to starting conditions in 2 min and re-equilibrating for 7 min, for a total run time of 120 min. The Fusion Lumos was operated in data-dependent mode, with both MS1 and MS2 scans acquired in the Orbitrap. The cycle time was set to 3 seconds. Monoisotopic Precursor Selection (MIPS) was set to Peptide. The full scan was done over a range of 400-1500 m/z, with a resolution of 120,000 at m/z of 200, an AGC target of 4e5, and a maximum injection time of 50 ms. Peptides with a charge state between 2-5 were picked for fragmentation. Precursor ions were fragmented by higher-energy collisional dissociation (HCD) using a collision energy of 36% and an isolation width of 0.7 m/z. MS2 scans were collected with a resolution of 50,000, a maximum injection time of 105 ms, and an AGC setting of 1e5. Dynamic exclusion was set to 45 seconds.

For Q Exactive Plus LC-MS/MS analysis, peptides were injected onto a homemade 30 cm C18 column with 1.8 um beads (Sepax), with an Easy nLC-1000 HPLC (Thermo Fisher), connected to a Q Exactive Plus mass spectrometer (Thermo Fisher). Solvent A was 0.1% formic acid in water, while solvent B was 0.1% formic acid in ACN. Ions were introduced to the mass spectrometer using a Nanospray Flex source operating at 2 kV. The gradient began at 6% B and held for 2 minutes, increased to 30% B over 85 min, increased to 50% B over 10 min, then ramped up to 70% B in 4 min and was held for 5 min, before returning to starting conditions in 4 min and re-equilibrating for 10 min, for a total run time of 120 min. The Q Exactive Plus was operated in data-dependent mode, with a full scan followed by 10 MS/MS scans. The full scan was done over a range of 400-1700 m/z, with a resolution of 70,000 at m/z of 200, an AGC target of 1e6, and a maximum injection time of 50 ms. Peptides with a charge state between 2-5 were picked for fragmentation. Precursor ions were fragmented by higher-energy collisional dissociation (HCD) using a collision energy of 35 and an isolation width of 1.0 m/z, with an offset of 0.3 m/z. MS2 scans were collected with a resolution of 35,000, a maximum injection time of 120 ms, and an AGC setting of 1e5. The fixed first mass for the MS2 scans was set to 120 m/z to ensure TMT reporter ions were always collected. Dynamic exclusion was set to 25 seconds.

#### Thermodynamic stability measurements of lysozyme

To measure intrinsic fluorescence, a solution of 10 μM chicken egg white lysozyme (Fisher) was prepared in 20 mM sodium phosphate buffer (pH 7.4) containing either 0 M or 7 M GdmCl. Denaturation was performed at the final concentrations of 0, 0.3, 0.6, 0.9, 1.2, 1.5, 1.8, 2.1, 2.4, 2.7, 3.0, 3.2, 3.4, 3.6, 3.8, 4.0, 4.2, 4.4, 4.6, 4.8, 5.0, 5.2, 5.4, 5.6, 5.8, 6.0, 6.2, 6.4, 6.6, 6.8, or 7.0 M GdmCl. Samples were then aliquoted out in triplicate in a black 96-well (Evergreen) microplate, incubating for 4 hr at RT. Samples were then analyzed on a SpectraMax M2 spectrophotometer, with a 280 nm excitation wavelength and emission wavelengths measured from 300-460 nm in steps of 5 nm. The relative fluorescence emission data at 360 nm was subsequently used to measure the fraction of unfolded protein.

For HR-SPROX analysis, lysozyme was dissolved at a concentration of 2 mg/mL in 20 mM sodium phosphate buffer (pH 7.4) and prepared using the method above, with the following modifications: Final GdmCl concentrations were 0, 0.3, 0.6, 0.9, 1.3, 1.6, 2, 2.4, 2.6, 2.8, 3, 3.2, 3.4, 3.6, 3.8, 4.0, 4.2, 4.4, 4.6, 4.8, 5, 5.5, 6, and 7 M. In place of fractionation, lysozyme samples were desalted using lab-made C18 spin columns. Columns were first conditioned using 50% ACN in 0.1% TFA, and 0.1% TFA. Samples were resuspended in 50 μL of 0.1% TFA and added to the columns. After washing the columns with 0.1% TFA, samples were eluted with 50% ACN in 0.1% TFA.

#### Database searches and quantitation

All MS2 data were searched against the *H. sapiens* UniProt database (20,975 entries, downloaded 6/18/2017) using the integrated Andromeda search engine with MaxQuant software(61). The searches included the TMT10-plex labels as fixed modifications and methionine oxidation as variable modifications and allowed for up to two missed cleavages with maximum false discovery rate (FDR) thresholds of 1%. For searches described in Figure S1, additional variable modifications were included as described. The complete parameter settings for MaxQuant searches and the search results are provided in Supplementary Tables 1-3.

#### Measurement of [GdmCl]_1/2_, ΔG_folding_, m-values and baseline oxidation levels

Subsequent analyses of MaxQuant search results and quantitations were conducted with scripts written in house using Mathematica (Wolfram). MaxQuant searches produced reporter ion intensities for 15 different TMT 10-plex experiments for -TMAO analyses and 6 for +TMAO experiments, with each tag corresponding to a different GdmCl concentration within a methionine oxidation denaturation curve. Reporter ion intensities within each 10-plex experiment were normalized with respect to one another using the median intensities of non-methionine-containing peptides. For each 10-plex experiment, the data for methionine-sulfoxide-containing (Met-O) peptides were converted to fraction oxidized by using the intensities of no-H_2_O_2_/0M GdmCl and highest concentration GdmCl data-points as initial and final baselines, respectively. For methionine-containing (Met) peptides, the inverse fractions were obtained by reversing the baselines. For sequences containing more than one methionine, the fully oxidized and fully unoxidized peptides were analyzed as Met-O and Met variants, respectively. Initially, the amplitudes and midpoints of denaturation curves ([GdmCl]_1/2_) for Met and Met-O containing peptides within each 10-plex experiments was measured by least squares fitting of each curve with the following sigmoidal equation:

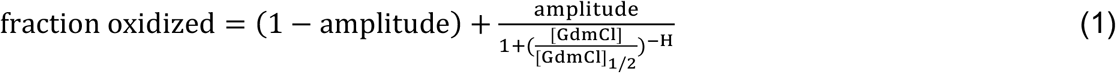

Where H is the largest absolute value of the slope of the curve. The amplitude and midpoints of peptides shared between different TMT 10-plex experiments were shown to be generally well correlated (Figure S2). The fraction oxidized data for each peptide contained in different TMT 10-plex experiments were merged by using the no-H_2_O_2_/0M GdmCl, H_2_O_2_/0M GdmCl and H_2_O_2_/highest GdmCl concentration as shared anchor points. The merged data (containing up to 300 individual [GdmCl] for each peptide (Figure S3)) were then fit to equation 12 (see below) by least squares fitting and [GdmCl]_1/2_, ΔG_folding_, m-values and baseline oxidation levels were determined. The two-state model is formally described below. Measured parameters for each peptide are provided in Supplementary Table 4.

#### Two-state model of methionine oxidation

We start by considering a two-state protein with a single methionine that can exist in completely folded or completely unfolded states:

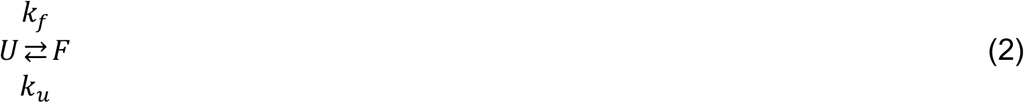

Where *k_f_* and *k_u_* are the first order rate constants for folding and unfolding, respectively. *K_folding_* is the folding equilibrium constant and is related to the folding and unfolding rate constants (k_f_ and k_u_) and thermodynamic stability (Δ*G_folding_*) by the following relationships:

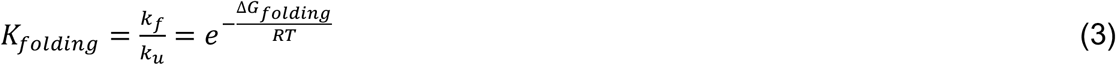

Where *R* is the gas constant, and *T* is the absolute temperature in Kelvin. Using the linear extrapolation model (LEM), the apparent free energy of folding changes linearly with respect to denaturant concentration ([D]):

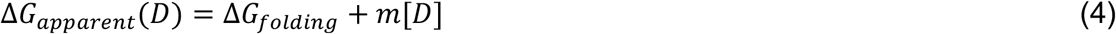

Where *ΔG_apparent_* is the free energy measured at a given concentration of denaturant, *ΔG_folding_* is the free energy in absence of denaturant, and *m* is the dependence of *ΔG_apparent_* on denaturant concentration measured as the slope of the *ΔG_apparent_* vs. *[D]* line. Thus, *K_folding_* at any given denaturant concentration can be described in terms of the apparent *ΔG_folding_*:

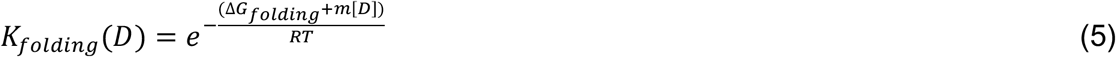

The rate of methionine oxidation will differ depending on whether the protein is in a folded or unfolded state. We refer to the pseudo-first order rate constants of oxidation within the folded or unfolded states as 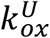 and 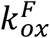, respectively. Thus, assuming first order kinetics, the fraction of the oxidized proteins within each state as a function of time can be defined as:

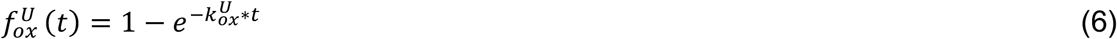

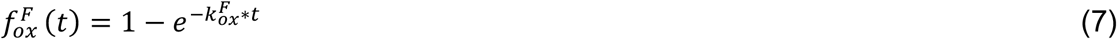

For a two-state protein, the conformational population will be divided into folded and unfolded fractions (*f_F_* and *f_U_*), in accordance to *K_folding_*, which is a function of *[D]* as shown in equation 4:

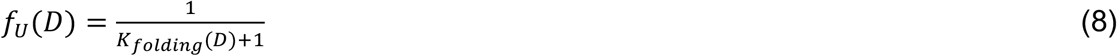

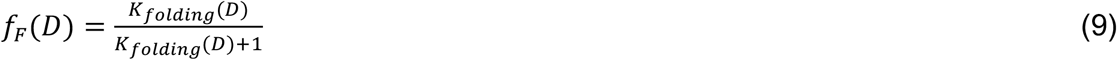

The overall fractional oxidation of the protein at a given time and denaturant concentration can be approximated by the following equation:

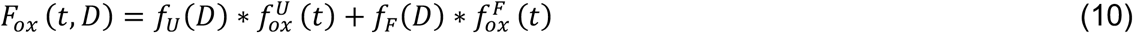

Note that equation 10 is making the assumption that H_2_O_2_ oxidation is occurring in a “pulse labeling” regime where folding and unfolding kinetics does not influence the rate of oxidation. For justification of this approximation refer to the next section. We normalize the overall fractional oxidation with respect to the initial and final baselines (when the protein is fully folded or unfolded, respectively):

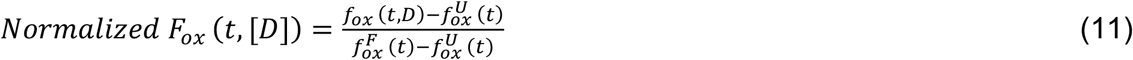

Substituting with equations 8 and 9, equation 11 simplifies to:

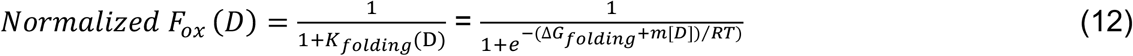

Note that while *F_ox_* is a function of denaturant concentration, time, and the rate constants for oxidation for both the folded and unfolded states, *Normalized F_ox_* is solely a function of *K_folding_ (and thus ΔG_folding_, m and [D]*) and is not dependent on time and the rate constants for oxidation. The denaturant concentration at the midpoint of transition *([D_]1/2_)* represents the point where *ΔG_apparent_* = 0 and is related to *ΔG_folding_* and *m* by the following equation:

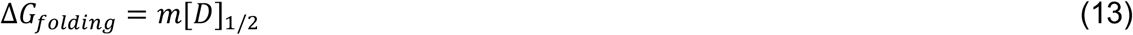

#### Justification of “pulse labeling” approximation

In the model presented in the previous section, we are making the simplifying approximation that methionine oxidation is occurring in a “pulse labeling” regime. In other words, we are assuming that the labeling time is not sufficiently long to allow for a significant fraction of the folded protein population to unfold prior to oxidation. This model differs from the “continuous labeling” model that is commonly used to interpret hydrogen-deuterium exchange (HDX) experiments (42) and was previously used by West et al. for analysis of SPROX data (20). In the “continuous labeling” model, it is assumed that localized unfolding occurs prior to oxidation. Below, we show that under our experimental conditions, the two models provide nearly identical *ΔG_folding_* measurements, justifying the use of the simpler “pulse labeling” model.

According to the classical HDX-like continuous labeling model (42, 62):

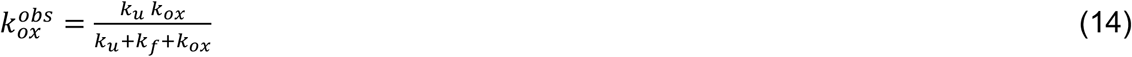

Where 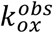 is the observed rate of oxidation, *k_f_* and *k_u_* are the rate constants for folding and unfolding, and *k_ox_* is the rate constant of methionine oxidation from the unfolded state. Under EX2 conditions where *k_f_* or *k_u_* are much greater than *k_ox_* (42):

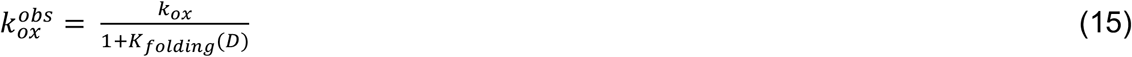

Where *K_folding_* is a function of denaturant concentration *([D])* as defined in equation 5. The observed fractional oxidation (*F_ox_*) as a function of time and *[D]* is defined as:

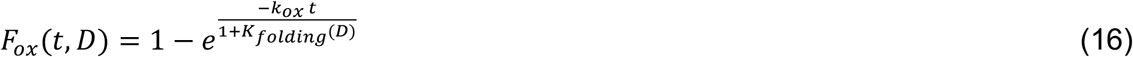

In our experiments, *F_ox_* is normalized with respect to the fully unfolded protein (where *K_folding_* = 0):

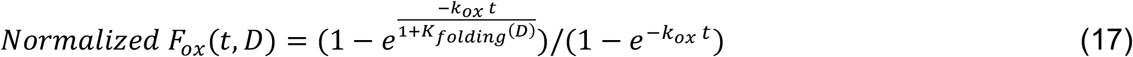

Note that whereas in the “pulse labeling” model, *Normalized F_ox_* is solely a function of *K_folding_*, in the “continuous labeling” model, it is additionally contingent on *k_ox_* and time. In this regime, the denaturant concentration at the midpoint of transition *([D]_1/2_*) does necessarily represent the point where *ΔG_apparent_* = 0 and is related to *ΔG_folding_* by the following equation (derived from equations 4, 5 and 17):

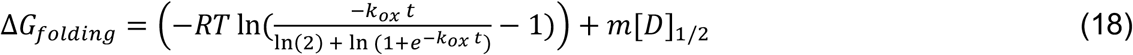

Comparing equations 13 and 18, we can state that 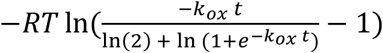 represents an offset between *ΔG_folding_* measurements obtained by the two models. As the term *k_ox_*t* approaches zero, this offset also approaches zero and the “pulse labeling” approximation becomes more valid. *k_ox_* is a pseudo-first order rate constant whose value is dependent on the concentration of H_2_O_2_ and the second order rate constant for methionine oxidation from the unfolded state. Although the latter is likely to be sequence dependent, we can use the rate of oxidation of free methionine as an approximation. This second order rate constant has been previously measured as 32.07 M^−1^h^−1^ (63). Thus, given our oxidation time (60 s) and H_2_O_2_ concentration (3% or 0.88 M):

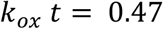

and:

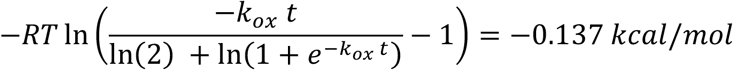

Thus, under our experimental conditions, *ΔG_folding_* measurement obtained using the simplifying “pulse labeling” approximation are within 0.14 kcal/mol of the “continuous labeling” model. As the conditions of the SPROX experiments deviate from this approximation (e.g. longer pulse times or higher H_2_O_2_ concentrations), the “pulse labeling” approximation will underestimate *ΔG_folding_* values to a greater extent.

#### Analysis of neighboring residue effects on baseline oxidation

Combined peptide data were binned based on the identities of the residues located at – 3, −2, −1, +1, +2 and +3 positions in the primary sequence relative to methionine. Sequences containing more than one methionine were excluded in this analysis.

Median baseline oxidation levels for each bin were measured, excluding far outliers (>2 SDs). Since tryptic peptides contain lysines and arginines at their carboxyl termini, −1, −2 and −3 bins for these two amino acids only contained a few mis-cleaved peptides and were excluded in this analysis.

#### Analysis of correlation between secondary structure and baseline oxidation

Mappings of UniProt accessions to protein data bank (PDB) structures were obtained from the Structure Integration with Function, Taxonomy and Sequence (SIFTS) database (downloaded August 2018) (64). DSSP secondary structure assignments were obtained from the PDB (downloaded August 2018). Using this data, 2,526 methionines in our dataset were mapped to DSSP secondary structures (Supplementary Table 5). For each secondary structure, the baseline oxidation levels for the corresponding methionines were collected. The selected peptides were limited to those for which more than two independent data points were collected for measurement of fractional oxidation. The distribution and median of the peptide measurements were analyzed, excluding outliers (>2 SDs). In cases were multiple structures and DSSP assignments were available for a given methionine, the measurements corresponding to that methionine were included for all matched secondary structures.

#### Analysis of correlation between solvent accessibility and baseline oxidation

UniProt accessions were mapped to PDB structures as above. For each mapped methionine, solvent accessible surface areas (SASA) were calculated from the associated PDB structures using the visualizing molecular dynamics (VMD) family of algorithms. For each structure, the coordinates of sulfur atoms belonging to methionines were found and analyzed for solvent accessibility using VMD’s *atomselect* and *sasa* commands, respectively. Solvent accessibility calculations were conducted using a probe radius of 1.4 angstroms and restricted to the space not occupied by another atom. A subset of PDB files were excluded from the analysis due to substitution of methionine with selenomethionine. In cases where multiple structures were mapped to a methionine, the median solvent accessible surface area was assigned to that methionine. Using this approach, we determined SASA values for 1,307 methionines in our dataset for which more than two independent data points were collected for measurement of fractional oxidation (Supplementary Table 5). Methionine baseline oxidation levels for specific ranges of SASA were collected and the distribution and median of the measurements were analyzed, excluding outliers (>2 SDs).

#### Assignment of intrinsically disordered proteins (IDPs)

A database of disordered protein regions was obtained from MobiDB3.0 (37). Sequences classified as “derived”, in which disorder was determined based on a number of different feature characteristics of the PDB structures (65), were collected and mapped to 2,896 methionines in our dataset for which more than two independent data points were collected for measurement of fractional oxidation (Supplementary Table 5). Assignments in MobiDB are divided into “disordered” and “conflict” based on whether a consensus regarding their annotation exists using different guidelines for assignment of disorder (37). The distribution and median of the baseline oxidation levels for methionines mapped to each category was determined, excluding outliers (>2 SDs).

#### Gene ontology (GO) analyses

Mappings of GO terms to UniProt accessions, and their relational hierarchies were obtained from the gene ontology database (downloaded June 2018) (66). For all proteins mapped to a given GO term, all methionine baseline oxidation levels, [GdmCl]_1/2_ and ΔG_folding_, values were collected. The p-values for differences between the distribution of these values for a given GO term, in comparison to the entire dataset, were determined using the Mann-Whitney U test. GO terms having p-values less than 10^−6^ for fractional oxidation and less than 10^−3^ for [GdmCl]_1/2_, m-value and ΔG_folding_ were identified for each parameter (Supplementary Table 6) and their connectivity was visualized using the program ReviGO (67) after excluding highly redundant terms (Figures S5–S6). For a selection of terms with the most significant p-values, the distribution and median of the parameters were compared after excluding outliers (>2 SDs) (Figures 4,5).

#### Analyses of intra-domain variability

Domain definitions and sequence boundaries were downloaded from the Pfam database (version 31.0, downloaded June 2018) (49). A total of 1,055 methionine-containing peptides for which [GdmCl]_1/2_,ΔG_folding_ and m-values were measured were mapped to 484 different Pfam domains (Supplementary Table 5). In analyzing the intradomain distribution and CVs of these three parameters (e.g. Figure 6), only domains with more than one mapped methionine were considered.

#### Data and software availability

All analyzed data and measurements can be found in supplemental data files. The raw and processed MS proteomics data have been deposited to the ProteomeXchange Consortium via the PRIDE partner repository with the dataset identifier PXD011456.

#### Supplementary Figures

**Figure S1.**
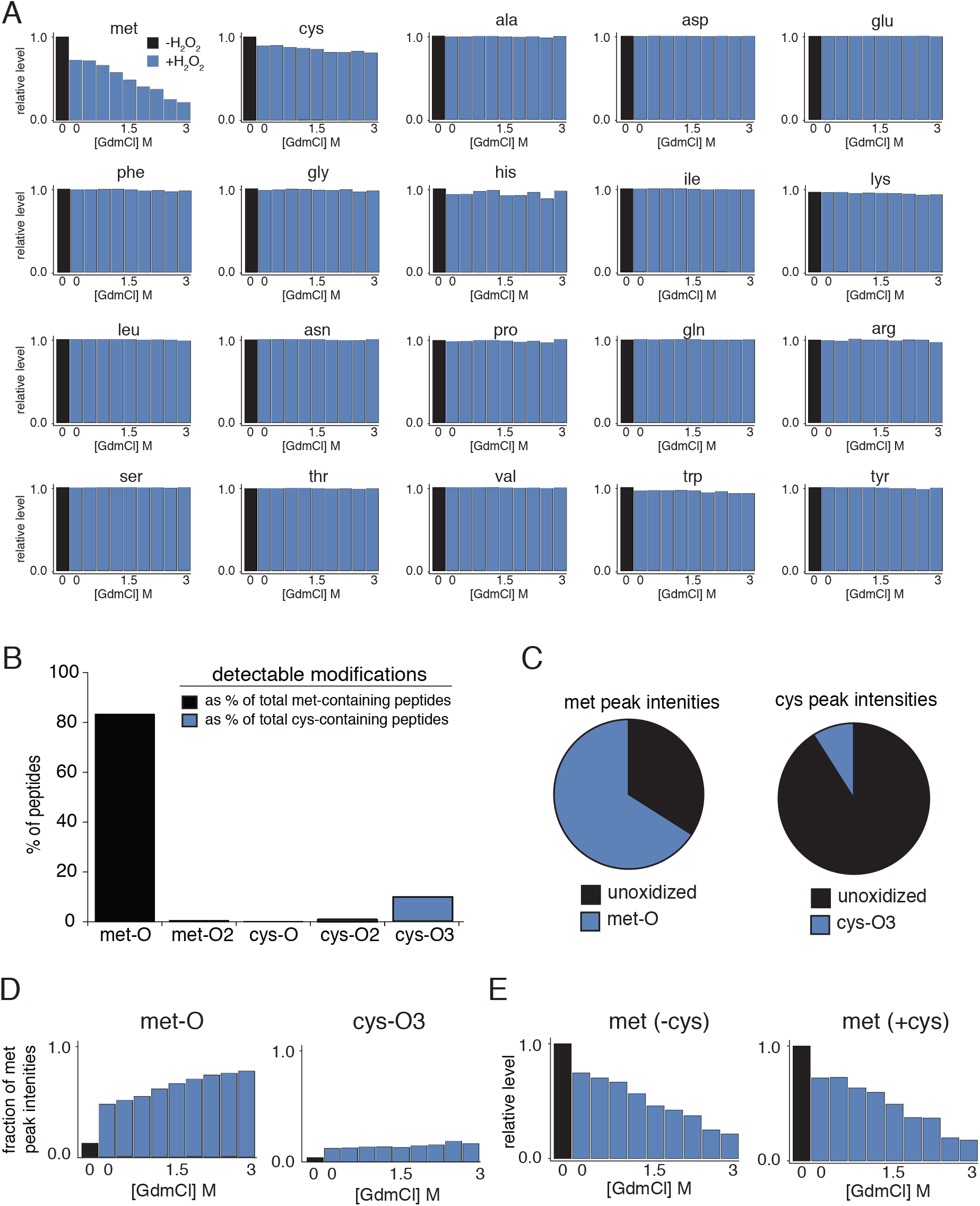
Proteome-wide prevalence of oxidized residues in HR-SPROX data. (A) Relative change in levels of all peptides containing unoxidized versions of each amino acid in the presence and absence of H_2_O_2_ and GdmCl. The y-axes indicate the median level of all peptides harboring one or more of the indicated residues relative to the level present in unoxidized samples. The data suggest that the oxidation of methionine (leading to decreases in levels of unoxidized methionines) is the most prevalent among all residues and increases as a function of [GdmCl]. The only other unoxidized residue that decreases significantly upon addition of H_2_O_2_ is cysteine. (B) To identify the oxidized forms of methionines and cystines present in our samples, we searched the proteomic data (technical replicate 1) for methionine sulfoxides (Met-O), methionine sulfones (Met-O_2_), sulfenic acid (Cys-O), sulfinic acid (Cys-O_2_), and sulfonic acid (Cys-O_3_). The bar plot indicates the fraction of methionine- and cysteine-containing peptide sequences in the dataset for which the indicated oxidized forms were also identified. (C) The fraction of the total intensities of the MS1 peaks of methionine and cysteine containing peptides that were mapped to oxidized forms of each peptide. The data shown in B-C indicate that Met-O and Cys-O_3_ are the only oxidized forms of methionine and cysteine that are present at significant levels. (D) The change in the relative levels of Met-O and Cys-O_3_ as a function of H_2_O_2_ and [GdmCl] as determined by the fraction of the total intensities of the corresponding MS2 reporter ion peaks mapped to oxidized forms of each peptide. The data indicate that unlike Met-O, Cys-O_3_ remains at low constant levels upon increasing GdmCl concentrations. This is likely due to the fact that most oxidizable cysteines are on protein surfaces and their exposure to oxidation is not influenced by denaturant-induced unfolding. (E) The disappearance of unoxidized met as a function of H_2_O_2_ and [GdmCl] for met-containing peptides that do or do not contain cystines. The data show that co-occurrence of cystines generally do not alter denaturation curves as monitored by met oxidation. Together, the data in this figure show that met-O is the dominant oxidized species formed in HR-SPROX experiments and although Cys-O_3_ is also formed at low levels, it does not artifactually influence the observed oxidation trends for methionine. Relevant to Figure 3.

**Figure S2.**
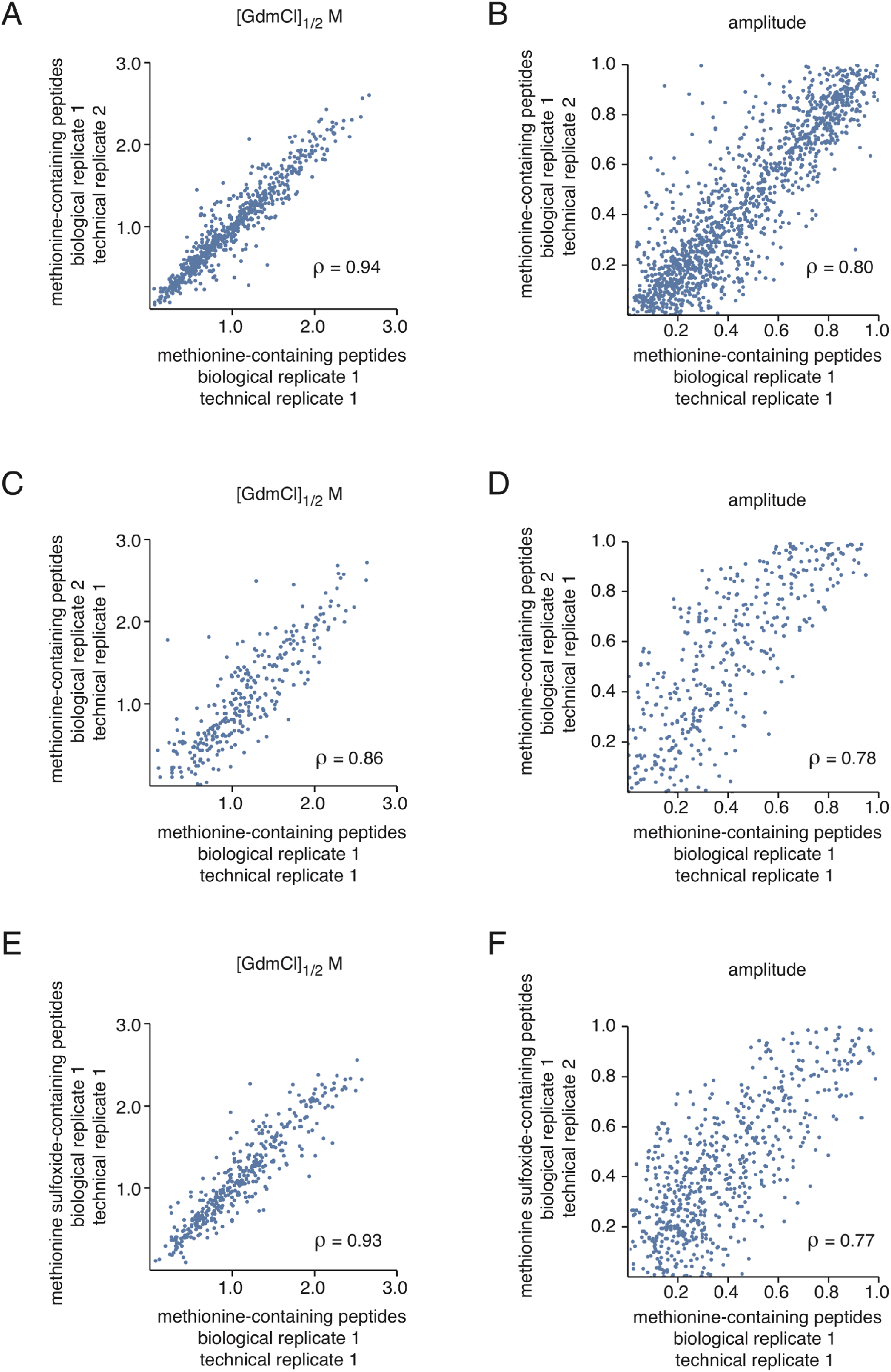
Pairwise comparisons of replicate HR-SPROX experiments and methionine versus methionine oxide-containing peptides. Data from each replicate experiment were fit to a model-independent sigmoidal equation (equation 1 in Supplementary Information) and amplitudes and [GdmCl]_1/2_ values were measured. The data were compared for pairs of technical (A-B) and biological (C-D) replicates, as well as for methionine and methionine sulfoxide-containing peptides (of the same sequence) within the same experiment (E-F). For each comparison, the Pearson coefficient (*ρ*) was measured. For all compared data (including comparisons of replicates not shown in this figure), the Pearson coefficient was greater than 0.7. Relevant to Figure 3.

**Figure S3.**
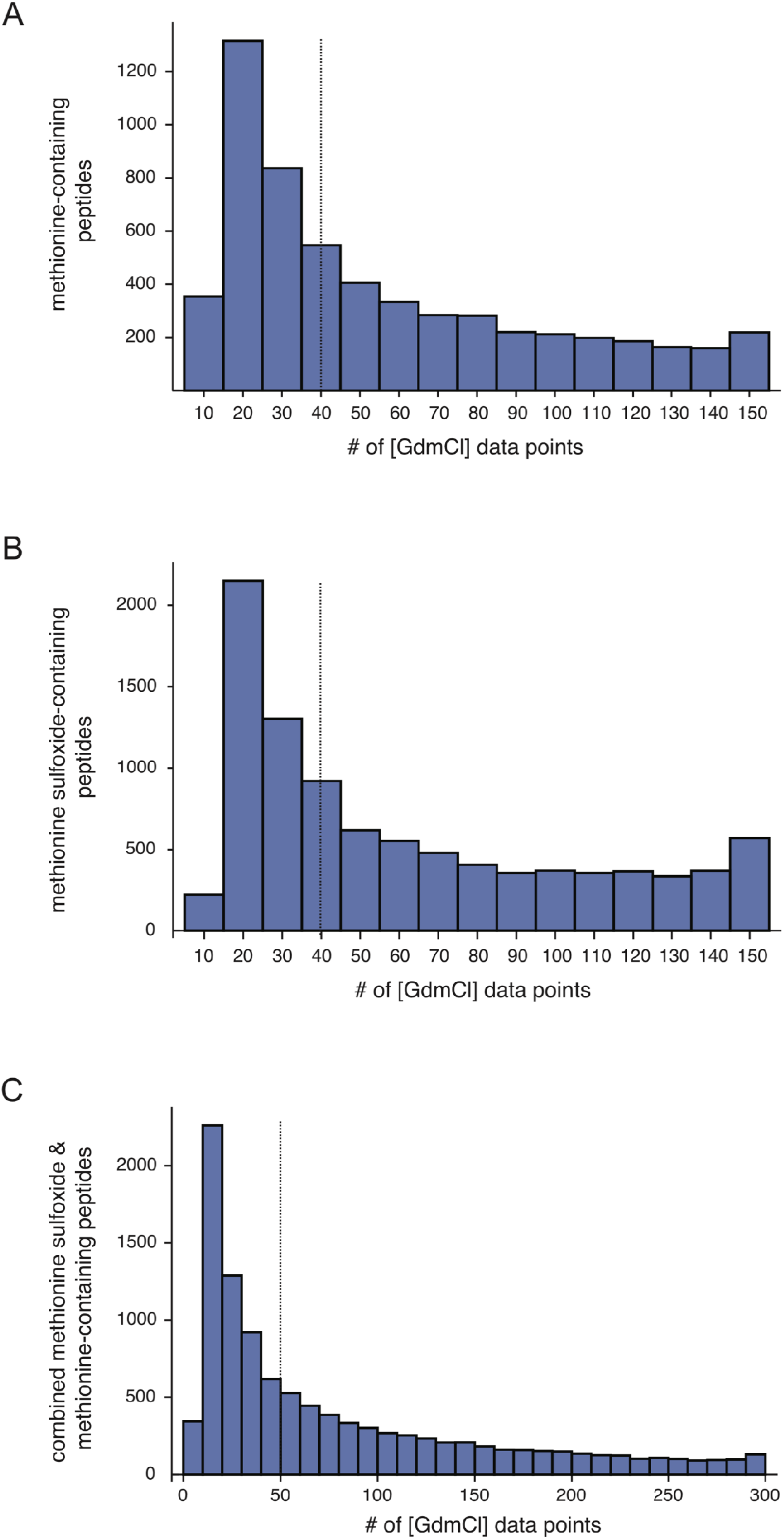
Distribution of the number of denaturation data-points used for HR-SPROX analysis of individual peptides. By combining 15 replicate experiments, we were able to obtain and analyze a large number of data-points (corresponding to different [GdmCl]) for each peptide in our dataset. The histograms indicate the distribution of the number of data-points used for the analysis of methionine-containing peptides (A), methionine sulfoxide-containing peptides (B) and merged data containing both methionine and methionine sulfoxide-containing peptides (C). The dotted lines indicate the median. Relevant to Figure 3.

**Figure S4.**
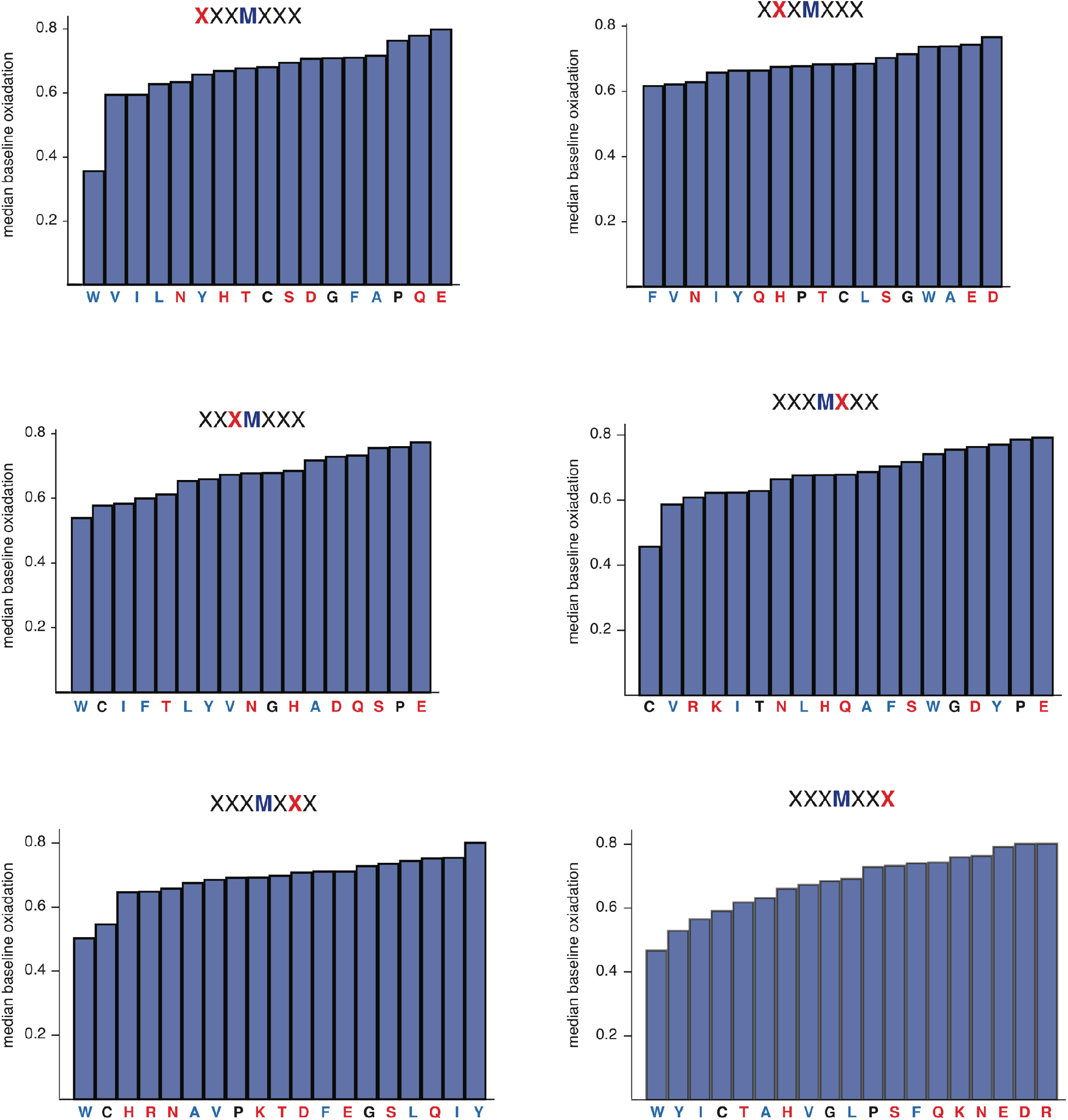
Median baseline methionine oxidation levels for peptides with varying methionine-neighboring residues. The bar plots indicate the median baseline oxidation levels for all methionine-containing peptides with specific residues at the specified position (red X) relative to the methionine in the primary sequence. Relevant to Figure 4.

**Figure S5.**
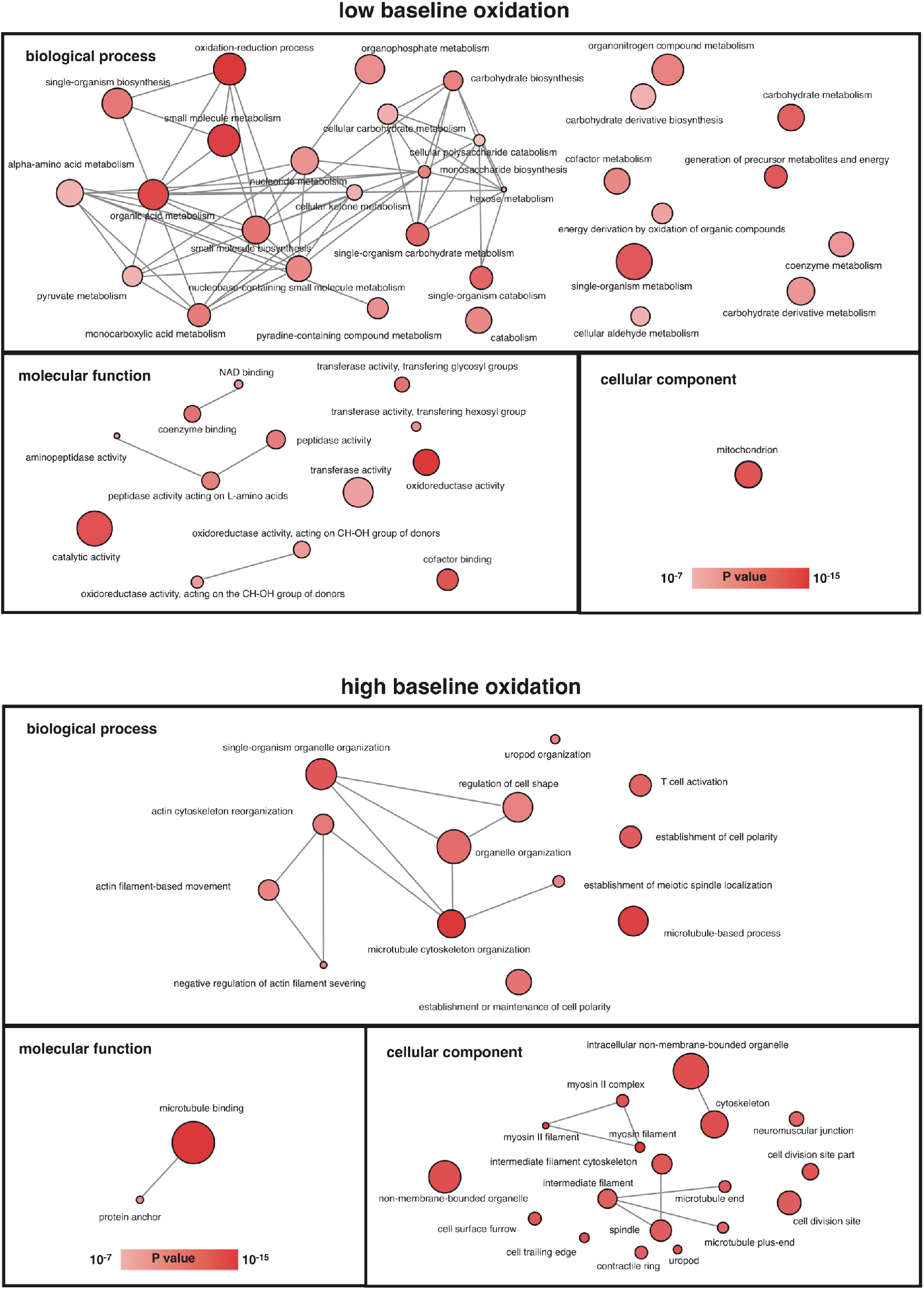
Gene ontologies (GO) enriched in proteins with high or low methionine baseline oxidation levels. GO terms enriched in proteins with high or low methionine baseline oxidation levels were identified and the p-value of the enrichment was calculated as described in Methods (Supplementary Table 6). The most significantly enriched GO terms were visualized by the software ReviGO to remove highly redundant terms and visualize the connectivity of the terms (gray lines). The size of the circles correlates with the number of proteins mapped to the GO term and the color indicates the p-value as shown in the bar scale. Relevant to Figure 4.

**Figure S6.**
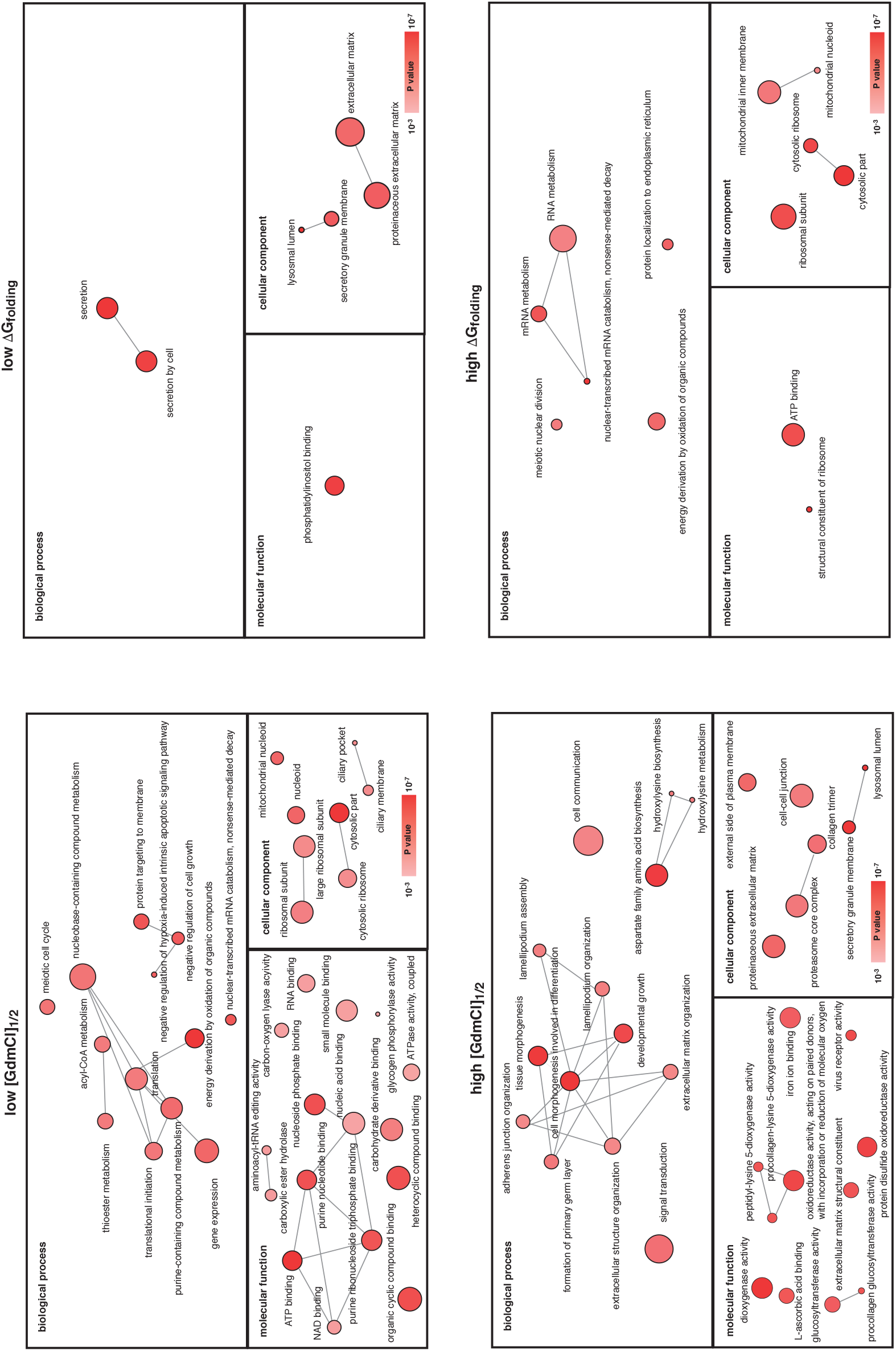
Gene ontologies (GO) enriched in proteins with high or low stabilities. GO terms enriched in proteins with high or low [GdmCl]_1/2_ or ΔG_folding_. Refer to Figure S5 for figure descriptions. Relevant to Figure 5.

**Figure S7.**
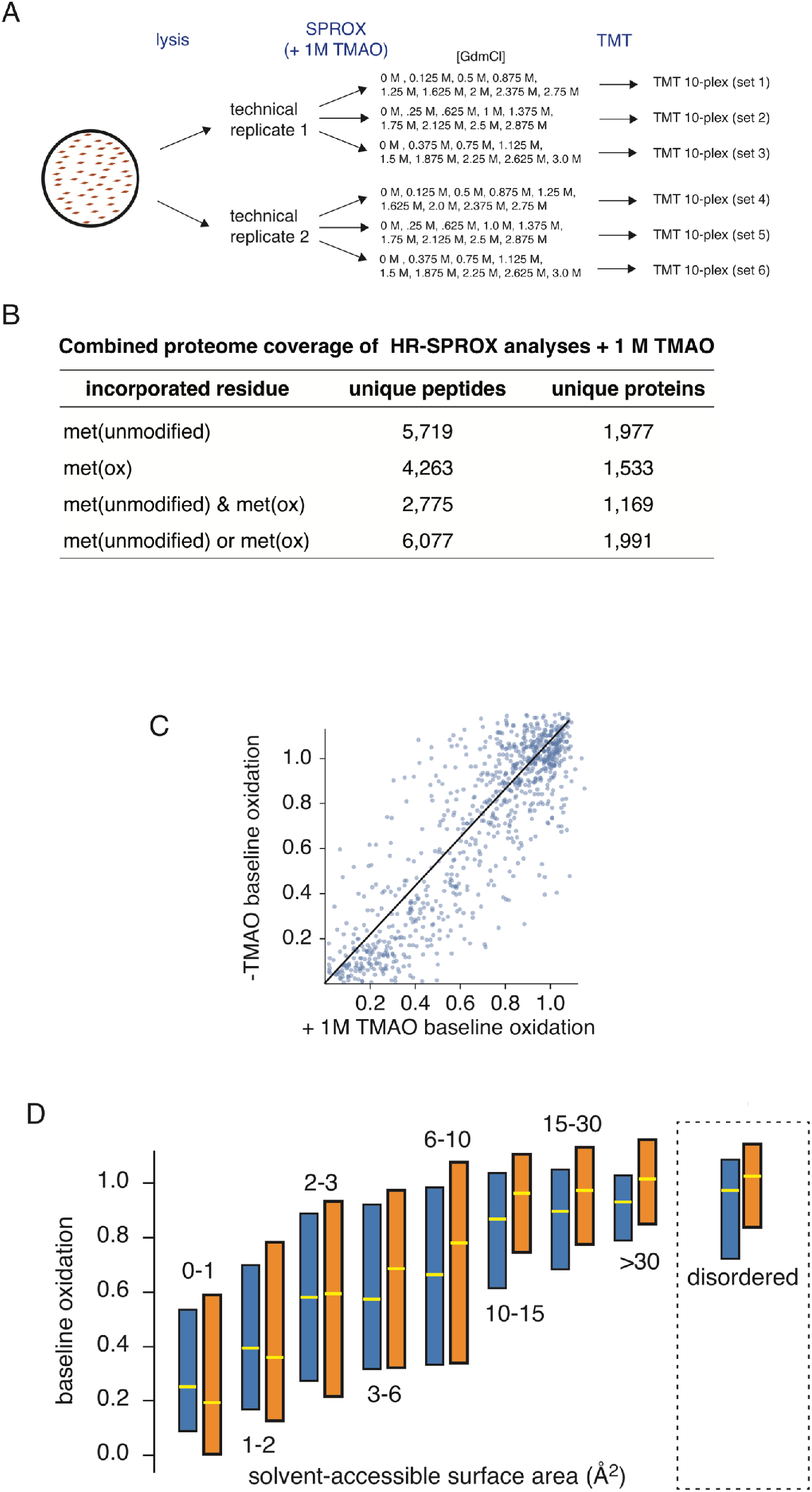
HR-SPROX analysis of the human fibroblast proteome in the presence of 1 M TMAO. (A) Schematic representation of the experimental design for HR-SPROX analysis of the human fibroblast proteome in the presence of 1 M TMAO, indicating the replicates and denaturant concentrations used in the study. Each technical replicate consists of 3 complementary sets of GdmCl concentrations tagged with TMT 10-plex reagents. (B) Coverage of proteomic analyses in the presence of 1 M TMAO. (C) Pairwise comparison of peptide baseline methionine oxidation levels with and without 1 M TMAO. Note that the oxidation levels of highly exposed methionines (top-right of the plot) are not impacted by TMAO. (D) Methionine baseline oxidation levels of highly exposed methionines, and those contained in regions classified as disordered in the MobiDB database, are not highly impacted by TMAO. Relevant to Figure 7.

#### Supplementary Tables

Supplementary Table 1. MaxQuant Search Parameters

Supplementary Table 2. Experiments and reporter ion definitions (+/− TMAO)

Supplementary Table 3. Peptide search results and reporter ion intensities for all experiments (+/− TMAO)

Supplementary Table 4. Measured folding parameters (+/− TMAO)

Supplementary Table 5. Term mappings for bioinformatic analyses (DSSP, SASA, disorder and Pfam)

Supplementary Table 6. Gene ontology (GO) enrichment data

AUTHOR CONTRIBUTIONS
E.W. and S.G. conceptualized the study, E.W., K.W., S.G. designed the experiments, E.W., K.W., J.H. performed the experiments, E.W., J.B., K.W., S.G. designed and carried out the data analysis, E.W. and S.G. wrote the manuscript.

